# Wide spectrum of neuronal and network phenotypes in human stem cell-derived excitatory neurons with Rett syndrome-associated *MECP2* mutations

**DOI:** 10.1101/2020.07.12.189621

**Authors:** Rebecca SF Mok, Wenbo Zhang, Taimoor I Sheikh, Kartik Pradeepan, Isabella R Fernandes, Leah C DeJong, Gabriel Benigno, Matthew R Hildebrandt, Marat Mufteev, Deivid C Rodrigues, Wei Wei, Alina Piekna, Jiajie Liu, Alysson R Muotri, John B Vincent, Lyle Muller, Julio Martinez-Trujillo, Michael W Salter, James Ellis

## Abstract

Rett syndrome (RTT) is a severe neurodevelopmental disorder primarily caused by heterozygous loss-of-function mutations in the X-linked gene *MECP2* that is a global transcriptional regulator. Mutations in the methyl-CpG binding domain (MBD) of MECP2 disrupt its interaction with methylated DNA. Here, we investigate the effect of *MECP2* L124W missense mutation in the MBD of an atypical RTT patient in comparison to severe *MECP2* null mutations. L124W protein had a limited ability to disrupt heterochromatic chromocenters due to decreased binding dynamics. We isolated two pairs of isogenic WT and L124W induced pluripotent stem cells. L124W induced excitatory neurons expressed stable protein, exhibited increased input resistance and decreased voltage-gated Na^+^ and K^+^ currents, and their neuronal dysmorphology was limited to decreased dendritic complexity. Three isogenic pairs of *MECP2* null neurons had the expected more extreme morphological and electrophysiological phenotypes. We examined development and maturation of L124W and *MECP2* null excitatory neural network activity using micro-electrode arrays. Relative to isogenic controls, L124W neurons had an increase in synchronous network burst frequency, in contrast to *MECP2* null neurons that suffered a significant decrease in synchronous network burst frequency and a transient extension of network burst duration. We capture these findings in a computational neural network model that shows the observed changes in network dynamics are best explained by changes in intrinsic adaptation currents in individual neurons. Our multilevel results demonstrate that RTT excitatory neurons show a wide spectrum of morphological, electrophysiological and circuitry phenotypes that are dependent on the severity of the *MECP2* mutation.

## INTRODUCTION

Rett syndrome (RTT) is a rare neurodevelopmental disorder affecting females that is primarily caused by heterozygous loss-of-function mutations in the gene *MECP2* (1). RTT is characterized by early developmental regression leading to microcephaly, loss of language and motor skills, development of hand stereotypies, gait abnormalities, and in some cases autism-like behaviours, autonomic dysfunction and seizures (2). *MECP2* is an X-linked gene that undergoes X chromosome inactivation (XCI) in female cells, resulting in mosaic expression of the wild-type (WT) and mutant *MECP2* alleles in RTT individuals that may alter symptom presentation in instances of non-random XCI (3, 4). Although *MECP2* mRNA is ubiquitously expressed, MECP2 protein is found to be most abundant in neurons (5), where it binds methylated DNA to act as a global transcriptional regulator to repress or activate genes (6, 7). A large number of RTT patient-associated *MECP2* missense mutations map to the Methyl Binding Domain (MBD) and disrupt the interaction with DNA required for proper MECP2 function (7).

Human stem cell-derived neuron models of RTT have confirmed many cellular phenotypes observed in *Mecp2* mouse models and RTT patients (8–15). In these studies, WT and *MECP2* point or null mutation ESCs and iPSCs were differentiated into Neural Progenitor Cells (NPCs) and then into electrically active cortical neurons. They showed that RTT neuron morphology is affected when MECP2 is dysfunctional, exhibiting reduced soma area, dendrite length, branch complexity and number of excitatory synapses. When RTT neuron electrophysiological properties were investigated during patch-clamp experiments (10, 12, 13), they show decreased action potential numbers in response to current pulses, fewer calcium transients and reduced frequency of spontaneous or miniature excitatory post-synaptic currents (mEPSC). These studies consistently demonstrated that RTT neurons have decreased size, dendritic branching, excitability and impaired excitatory neurotransmission, together suggesting a neuronal maturation defect.

It has been shown in electrophysiological studies of mouse cortical slices and hippocampal cultures that *Mecp2* neurons have decreased mEPSC amplitude or frequency (16, 17). This implies that decreased excitatory neuronal activity and alteration of activity-dependent gene expression regulated by MECP2 (18, 19) may alter circuit maturation dynamics and contribute to the observed RTT phenotypes. How these changes translate into patterns of activity across networks of neurons in the developing human brain remains poorly documented. Recent approaches using induced excitatory neurons cocultured with astrocytes on micro-electrode arrays (MEAs) now allow evaluation of the development of synchronous network activity and how that circuitry is affected in neurodevelopmental disorders (20–22). It is possible that early activity patterns of synchrony in neuronal networks observed during early brain development are impacted by the identified changes in single neurons, and that the degree of the impact depends on the severity of *MECP2* mutations. Simulation models predicting in vivo alterations in circuitry have advanced rapidly and provide an exciting opportunity to predict human circuitry changes in vitro such as variations of synchronic activity patterns during development (23).

In this work, we use a multilevel approach (molecular-cellular-network-modeling) to investigate the impact of a novel atypical RTT patient-associated missense mutation in the MECP2 MBD at L124W (c.371T>G). This variant has unknown molecular consequences on MECP2 heterochromatin binding. We compared atypical L124W to severely affected *MECP2* null excitatory neurons and their respective isogenic controls. We investigated protein expression levels, single neuron structure and electrophysiological characteristics (e.g., input resistance, voltage-gated Na^+^ and K^+^ currents, action potential firing, and excitatory synaptic function), and neuronal network dynamics using MEA recordings. The results were integrated into a computational network model of spiking neurons. Our results reveal a range of RTT-associated heterochromatin binding, neuron morphology, electrophysiology and circuitry phenotypes depending on *MECP2* mutation severity. The computational model simulations further indicate that the observed changes in RTT network dynamics are likely attributable to changes in intrinsic adaptation currents in single neurons.

## RESULTS

### Limited heterochromatin disruption by MECP2 L124W protein

MECP2 is known to play a role in chromatin architecture (24). The structure of crystallized human WT MECP2 MBD bound to the *BDNF* promoter (25) demonstrated that the location of L124 maps in close 3D proximity to R106 (Figure 1A). According to RettBASE (26), a database of patient *MECP2* variants, both of these residues are mutated in classical Rett syndrome patients to L124F and R106W, respectively. When these two variants were expressed in mouse cells they disrupted heterochromatic chromocenters (27–29). To investigate whether the novel L124W MBD missense mutation disrupts heterochromatin similarly to L124F and R106W, we assessed its ability to bind and cluster chromocenters relative to WT. In brief, we over-expressed MECP2-GFP fusions in C2C12 mouse myoblast cells, which have low levels of endogenous Mecp2. GFP colocalization with DAPI marked chromocenters was measured by confocal microscopy. WT showed distinctly formed chromocenter clusters with high colocalization of GFP and DAPI as expected, while R106W and L124F showed completely disrupted chromocenter clustering and indistinct GFP and DAPI colocalization (Figure 1B). In contrast, the novel L124W mutation showed limited heterochromatin disruption characterized by unaffected chromocenter number and reduced chromocenter size (Figure 1B,C). L124W also was more distributed through the nucleus than was WT, showing reduced colocalization of GFP with DAPI as measured by Pearson’s correlation coefficient (PCC) (Figure 1B,C).

**Figure 1.**
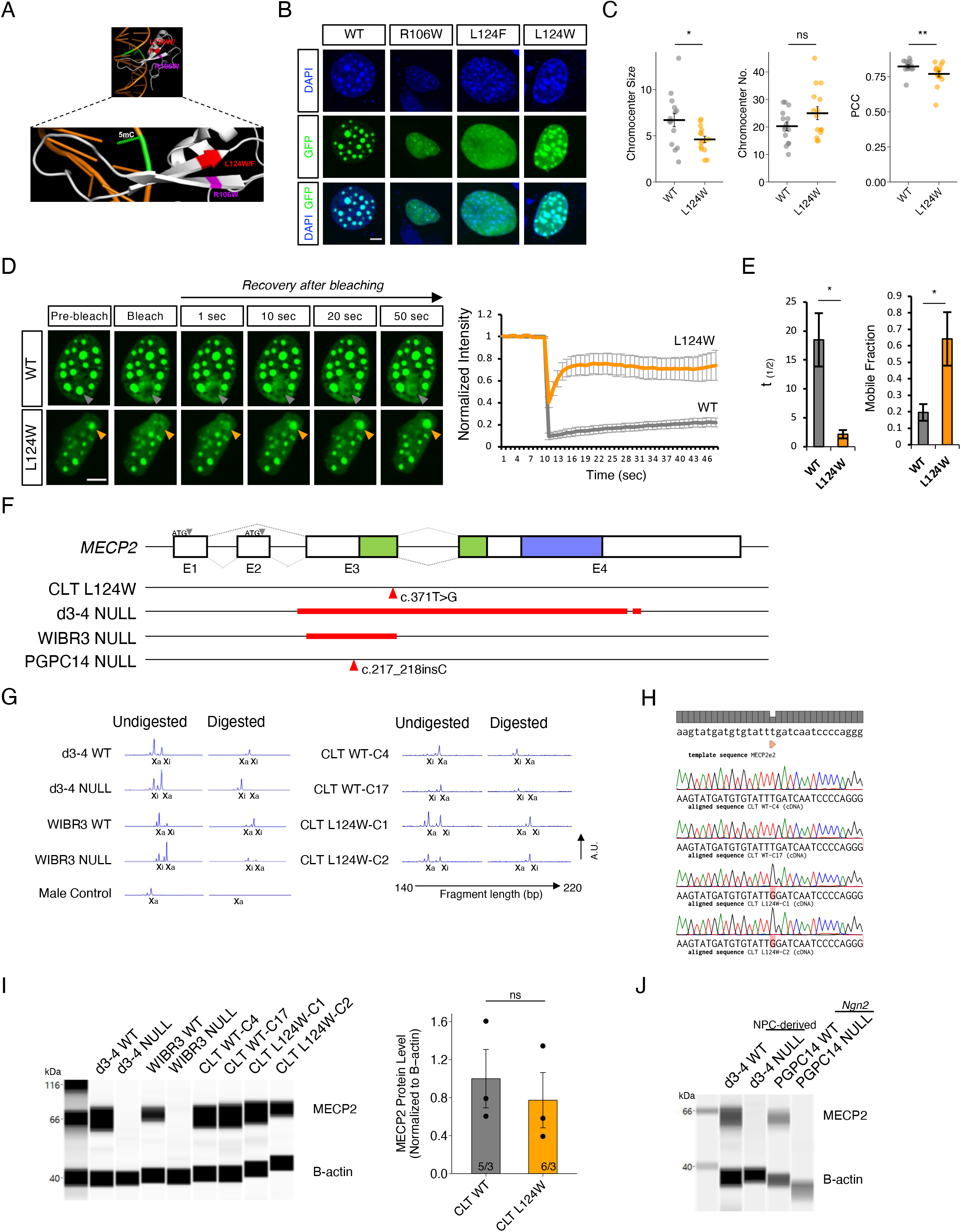
Molecular impact of MECP2 L124W missense mutation and generation of iPSC lines. **A.** MECP2 MBD 3D structure modeling L124W point mutation position relative to R106W. **B.** Representative images of C2C12 mouse myoblast cells overexpressing WT-GFP and mutant MECP2-GFP fusion proteins stained with DAPI (blue) and GFP (green). Scale bar = 2 μm. **C.** Quantification of chromocenter size and number and overlap by Pearson’s correlation coefficient (PCC). n = 15. **D.** Representative images of FRAP time course for GFP and quantified fluorescence intensity of WT-GFP and L124W-GFP in C2C12 mouse cells. Scale bar = 2 μm. **E.** Quantification of WT-GFP and L124W-GFP half-life and mobile fraction. n = 5. **F.** Schematic showing position of *MECP2* mutations in the iPSC/ESC lines used in this study. Red line indicates location of deletion and red arrow indicates location of point or frameshift mutation. **G.** Amplified peaks from PCR of polymorphic androgen receptor locus on the X active (Xa) or X inactive (Xi) for endonuclease digested and undigested gDNA samples from WT and *MECP2* mutant iPSC lines. **H.** Sanger sequencing of cDNA for the CLT WT or c.371T>G (p.L124W) *MECP2* mutation from restricted expression of the active X chromosome in iPSC clones. **I.** Digitized blot of isogenic *Ngn2* neurons and quantification of MECP2 protein level by capillary western in isogenic L124W *Ngn2* neurons normalized to β-actin. **J.** Wes capillary-based protein detection of MECP2 protein in 6 week old PGPC14 WT and null *Ngn2* neurons, compared to 4 week old d3-4 WT and null NPC-derived neurons. All data were shown as mean +/- SEM, indicating n/N = number of samples/number of replicate batches. Statistical significance was evaluated by Student’s t-test or Mann Whitney test, as appropriate, in Figure C, E and I. ns p > 0.05, *p < 0.05, **p < 0.01.

To explore the binding dynamics of L124W protein, fluorescence recovery after photobleaching (FRAP) live-imaging was used to study MECP2 protein mobility (29). WT and L124W MECP2-GFP fusion proteins were overexpressed in C2C12 cells, and following photobleaching of a single chromocenter focus, MECP2-GFP fusion protein signal recovered more rapidly in the L124W compared with the WT over time (Figure 1D). L124W had a shorter half-life and larger available unbound mobile fraction to recover the bleached signal that suggests it has decreased binding dynamics with heterochromatin (Figure 1E). Taken together with the partial impairments in chromocenter clustering, these results point towards a limited effect of L124W relative to L124F and R106W mutations on MECP2 chromocenter association due to decreased binding dynamics.

### Generation of MECP2 L124W iPSCs and *Ngn2*-derived excitatory neurons

As differences in genetic background and modifier genes contribute to RTT phenotype severity (30, 31), disease modeling strategies using isogenic WT cells provide the most precise controls for comparing to affected cells. By taking advantage of XCI in female somatic cells, we and others have generated isogenic pairs of RTT stem cell lines. As described below, we first generated isogenic human iPSC bearing the L124W missense mutation. As controls, we chose *MECP2* null cells including isogenic lines derived from a d3-4 patient (9) and a *MECP2* null line derived by gene editing of the human ESC line WIBR3 (12) (Figure 1F). We also generated a third *MECP2* null isogenic pair using gene editing of the PGPC14 healthy control iPSC line (32).

We generated iPSCs from the atypical RTT patient (CLT) harbouring a heterozygous *MECP2* c.371T>G variant resulting in the L124W amino acid change. iPSCs were reprogrammed from fibroblasts using Sendai virus. Expanded clones were assessed for XCI status using the androgen receptor (AR) endonuclease digestion assay to isolate isogenic lines fully skewed to either the WT or L124W allele on the active X chromosome (X_a_) with a male line as a control for full digestion of an unmethylated X_a_ (Figure 1G). The terminology used for these CLT isogenic cell lines is CLT-WT and CLT-L124W. Two lines of X_a_ CLT-WT (C4 and C17) and two lines of X_a_ CLT-L124W (C1 and C2) were obtained with normal 46XX karyotype (Figure S1A). Restricted mRNA expression of the X_a_ allele was confirmed by cDNA sequencing across the L124W variant (Figure 1H) and shown to not be subject to erosion of the inactive X chromosome (X_i_) over extended passage (33). The iPSC lines demonstrated pluripotency by positive staining for OCT4, NANOG, SSEA-4 and TRA-1-60 (Figure S1B), spontaneous differentiation into the 3-germ layers (Figure S1C) and pluripotent cell expression profiles determined by RNA-seq and Pluritest analysis (34) (Figure S1D).

To examine L124W protein levels in excitatory cortical neurons we used a rapid induced *Neurogenin-2* (*Ngn2*) overexpression protocol (35, 36) (Figure S1E). In brief, WT and RTT iPSCs/ESCs were transduced with the Tet-inducible *Ngn2-puro* lentivirus vectors, overexpression was induced with doxycycline for 8 days and selected with puromycin, and dividing cells were eliminated with Ara-C prior to excitatory neuron re-plating and maturation. As expected, MECP2 capillary-based protein detection of 6 week old *Ngn2* neuron extracts demonstrated absence of MECP2 protein in d3-4 and WIBR null lines, but the levels of MECP2 protein were similar in isogenic CLT-WT and CLT-L124W lines (Figure 1I). We conclude that L124W mutant protein is present and produced at unchanged levels in CLT-L124W cells.

While we proceeded to evaluate the CLT, d3-4 and WIBR3 neurons for neuronal activity as described in the next section, we used CRISPR/Cas9 gene editing to isolate a novel *MECP2* null line. We introduced indels into a previously characterized healthy control female PGPC14 iPSC line from the Personal Genome Project Canada (32). PGPC14 iPSCs were co-transfected with Cas9 and a gRNA specific for exon 3 of *MECP2*. Selected clones were screened for normal karyotype (Figure S1F) and found to be positive for hallmark pluripotency markers (Figure S1G). cDNA sequencing identified a 1 bp insertion (c.217_218insC) into *MECP2* exon 3 on the X_a_ (Figure S1H) of the PGPC14 null line, resulting in a frameshift and premature truncation of the protein (p.Val74Cysfs*16) before the MBD (Figure 1F). Generation of *Ngn2* neurons and protein detection by immunocytochemistry (Figure S1I), capillary (Figure 1J) and conventional western blot (Figure S1J) demonstrated that PGPC14 indel neurons were *MECP2* null, compared with d3-4 null samples generated from NPC derived neurons. The PGPC14 parental cells exhibit robust differentiation into many cell types, in contrast to the d3-4 lines, and were employed for subsequent neuron morphology and circuitry experiments.

### RTT *Ngn2* neurons exhibit a range of alterations in intrinsic membrane properties and excitatory synaptic function

To validate *Ngn2*-derived excitatory neurons as an appropriate cellular model of RTT, we examined their electrophysiological characteristics and first determined whether there were detectable differences in the known *MECP2* null associated phenotypes. We made whole-cell patch-clamp recordings to investigate the intrinsic membrane properties of 4-5 week-old *Ngn2* neurons from isogenic d3-4 and WIBR3 *MECP2* null lines co-cultured with mouse astrocytes. We looked for alterations in electrophysiological characteristics as previously reported in mouse iPSC-derived neurons and human RTT neurons conventionally differentiated via a neural precursor cell (NPC) stage (10, 37). Indeed, relative to isogenic controls *MECP2* null neurons exhibited depolarized resting membrane potential, higher input resistance and decreased cell capacitance (Figure 2A-C), consistent with the previous findings on RTT iPSC-derived neurons. We next assessed whether *MECP2* null mutant neurons had alterations in active membrane properties, as previously reported in RTT neurons (10, 37). Voltage-gated Na^+^ and K^+^ currents, under voltage-clamp condition, were evoked by depolarizing voltage steps from a standardized holding potential of −70 mV. We found that both of Na^+^ and K^+^ currents were diminished in d3-4 and WIBR3 null neurons, compared with their respective isogenic WT (Figure 2D). Given these alterations in the electrophysiological characteristics, *MECP2* null neurons were anticipated to have dysfunction in action potential firing. We therefore performed whole-cell current-clamp recordings to investigate the characteristics of action potentials, which were elicited by recording in current-clamp mode and injecting a series of current steps from −5 pA to 100 pA in both WT and *MECP2* null neurons (Figure 2E & S2A-E). The majority of action potential parameters were not significantly different across the groups (Figure S2A-E), dysfunction in the generation of action potentials was observed in *MECP2* null neurons as compared with the respective WT controls (Figure 2E). Together, these findings indicate that loss of *MECP2* causes alterations in electrophysiological characteristics and deficiency in action potential firing, which may lead to a further influence on the functions of the neuronal network. We next examined synaptic transmission by recording mEPSCs in 6-week-old *Ngn2* WT and *MECP2* null neurons from d3-4 and WIBR3. We observed reduced AMPAR-mEPSC amplitude and frequency in the *MECP2* null neurons (Figure 2F), consistent with previous reports of hypo-activity in RTT iPSC-derived cortical neurons (10, 37). Taken together, these data indicate that *MECP2* null neurons recapitulated known RTT-associated excitatory synaptic phenotypes.

**Figure 2.**
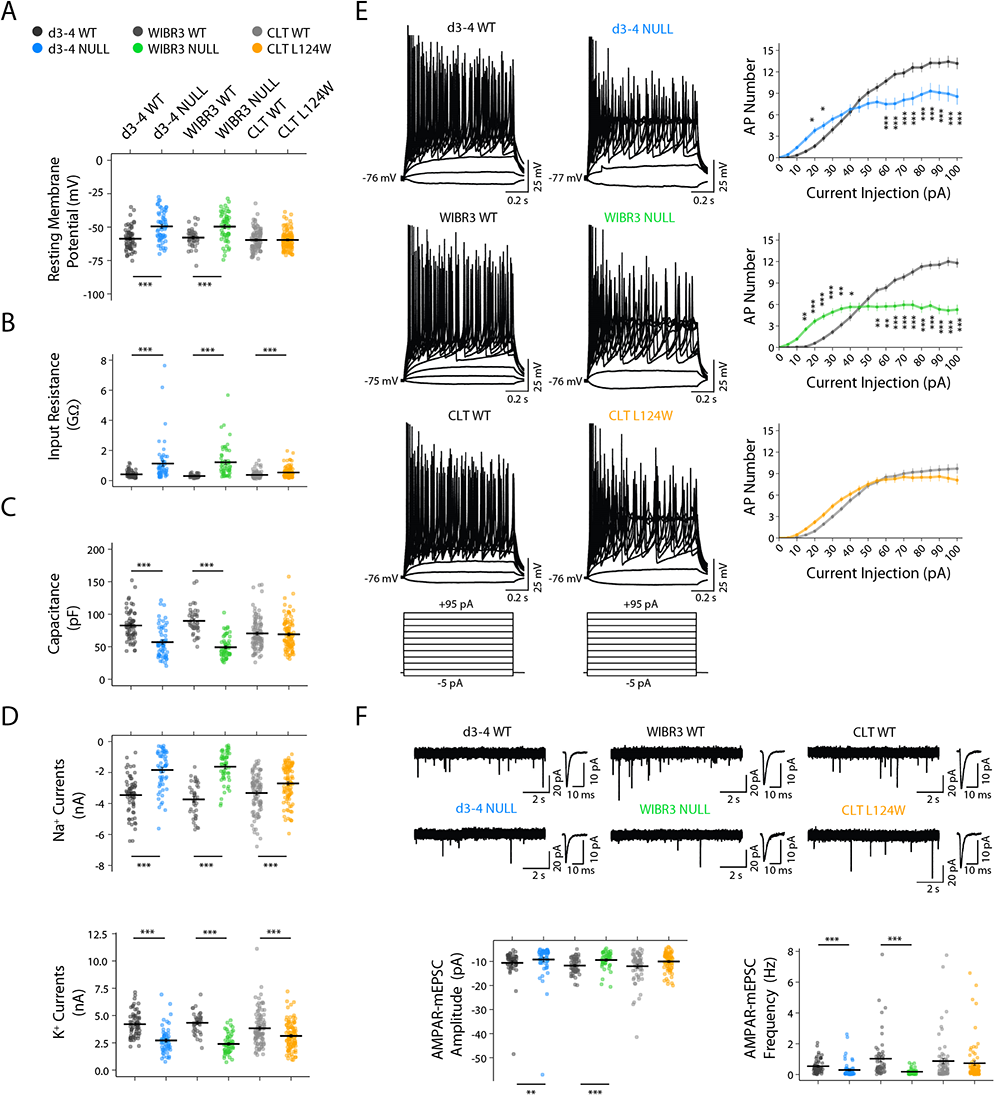
Deficits in intrinsic membrane properties and excitatory synaptic function in RTT neurons. (**A-D**) Scatter plots showing all data points for the resting membrane potentials (**A**), input resistance (**B**), cell capacitance (**C**), voltage-gated Na^+^ currents measured at −10 mV (**D**, Top), and voltage-gated K^+^ currents measured at +60 mV (**D**, Bottom) in 4-5 week-old Ngn2 neurons of WT ( n = 66, 42, and 99, respectively) and RTT from d3-4 (n =58), WIBR (n = 55), CLT (n = 100). The graphs also display average resting membrane potentials, input resistance, cell capacitance, Na^+^ currents and K^+^ currents. **E** Left, Representative traces showing evoked action potentials in 4-5 week-old Ngn2 neurons of WT and RTT from d3-4, WIBR and CLT. Right, Plots depict the numbers of evoked action potentials triggered by a series of current step from +5 pA to +100 pA for 1 s in 4-5 week-old Ngn2 neurons of WT (n = 66, 42, and 99, respectively) and RTT from d3-4 (n = 58), WIBR (n = 55), CLT (n = 100). **F**. Top, Typical traces display mEPSCs in 6 weeks old Ngn2 neurons of WT and RTT from d3-4, WIBR and CLT. Inset shows averaged mEPSCs in the neuron on the left. Bottom, Scatter plots display all data points for the amplitude and frequency of mEPSCs. The graphs also display average amplitude and frequency of mEPSCs in 6 weeks old Ngn2 neurons of WT (n = 56, 58, and 68, respectively) and RTT from d3-4 (n = 62), WIBR (n = 44) and CLT (n = 78). All data were shown as mean +/- SEM. Statistical significance was evaluated by Student’s *t*-test or Mann Whitney test, as appropriate, in Figure A-D and F, and by two-way repeated measure ANOVA followed by post hoc test in Figure E. *p < 0.05, **p < 0.01, ***p < 0.001.

Based on the limited disruption of heterochromatin binding by L124W MECP2 protein, we predicted that CLT L124W *Ngn2* neurons would have a milder cellular phenotype in comparison to their isogenic WT neurons. In CLT L124W neurons the input resistance was greater than in the CLT WT neurons. However, neither resting membrane potential nor cell capacitance in CLT L124W neurons were different from those in CLT WT neurons (Figure 2A-C). Voltage-gated Na^+^ and K^+^ currents in CLT L124W neurons were smaller than those currents in CLT WT neurons (Figure 2D). However, CLT L124W did not display deficiency in firing action potentials (Figure 2E). Next, when examining excitatory neurotransmission in CLT WT neurons and CLT L124W *Ngn2* neurons, neither AMPAR-mEPSC amplitude nor frequency in CLT L124W neurons were different from CLT WT neurons (Figure 2F). Therefore, recordings from CLT WT and CLT L124W *Ngn2* neurons show increased input resistance and impaired voltage-gated Na^+^ and K^+^ currents in the mutant neurons. These differences were also observed in *MECP2* null neurons which additionally showed differences, in comparison with their isogenic controls, in resting membrane potentials, cell capacitance, action potential firing, and excitatory synaptic function.

### Altered neuronal size, complexity and excitatory synapse density in RTT *Ngn2* neurons

Reduced size and complexity have been identified as prominent morphometric features of RTT using human NPC-derived cortical neurons (9, 12, 13). To assess morphometry in *Ngn2* neurons, 6 week old neuron co-cultures with mouse astrocytes were sparsely labelled with GFP in 3-5 independent experiments. Fixed cells were stained for DAPI, GFP and MAP2, and images captured on a confocal microscope reveal similar cell densities for all genotypes (Figure 3A). Soma area and dendrite tracing was performed by two observers blinded to genotype. As anticipated, *MECP2* null (d3-4, WIBR and PGPC14) neurons had decreased average soma area and total dendrite length, recapitulating the known RTT-associated phenotypes (Figure 3B,C). In contrast, the CLT L124W missense neurons exhibited no changes in soma area or dendrite length (Figure 3B,C).

**Figure 3.**
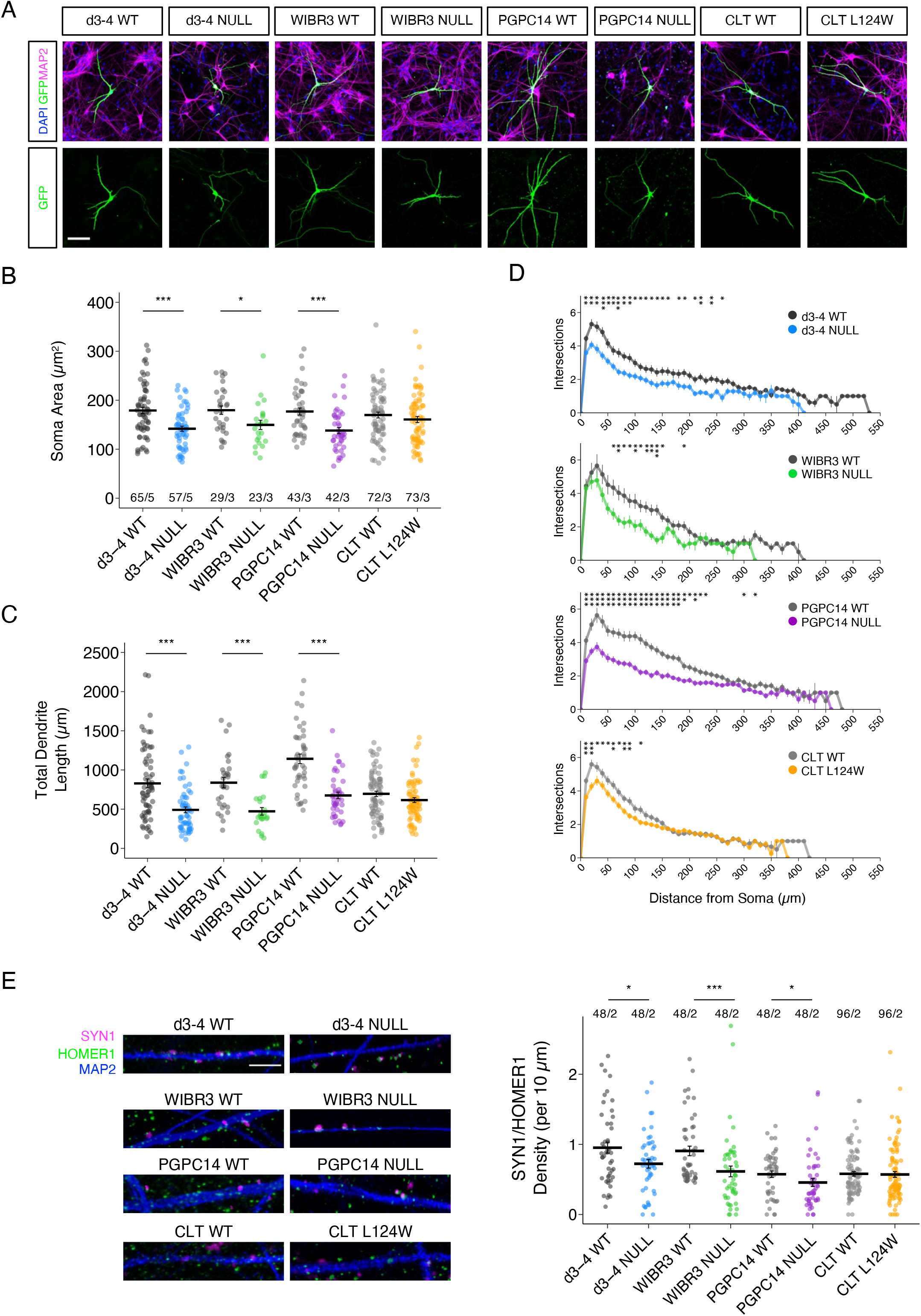
RTT Ngn2-derived neurons have reduced size and complexity. **A.** Representative images of 6 week old WT and RTT Ngn2 neurons co-cultured with mouse astrocytes, with a GFP sparse labelling (green), pan-neuronal marker MAP2 (magenta) and DAPI (blue). Scale bar = 100 µm. **B.** Neurons quantified for soma area, **C.** Total dendrite length, and **D.** Sholl complexity of individual neurons (10 µm increments from the soma). **E.** Representative images of 6 week old WT and RTT Ngn2 neurons co-cultured with mouse astrocytes, labelled with pre- and post-synaptic markers SYN1 (magenta) and HOMER1 (green), quantified for synapse density along dendrites labelled by MAP2 (blue). Scale bar = 5 µm. All data were shown as mean +/- SEM in Figure B-D, indicating n/N = total number of neurons/number of replicate batches, and in Figure E indicating n/N = total number of dendrite segments/number of replicate batches. Statistical significance was evaluated by Student’s t-test or Mann Whitney test, as appropriate, in Figure B-E. *p < 0.05, ** p < 0.001, ***p < 0.001.

However, Sholl analysis of the CLT L124W neurons revealed decreased branching complexity relative to WT, similar to the reduced branching complexity in *MECP2* null neurons relative to WT (Figure 3D & S3A). To investigate changes in synapses, excitatory synapse density was quantified based on co-localization of pre-synaptic SYN1 and post-synaptic HOMER1 puncta along dendritic MAP2 signal. The d3-4, WIBR3 and PGPC14 *MECP2* null neurons showed reduced SYN1/HOMER1 density compared to its respective isogenic control. On the other hand, CLT L124W neurons did not show significant changes relative to WT (Fig. 3E). Taken together, CLT L124W neurons show changes that are less pronounced than the ones in *MECP2* null neurons, consistent with more limited mutation-dependent phenotypes found in L124W protein heterochromatin association and excitatory neuron activity.

### Aberrant network circuitry in RTT *Ngn2* neuronal cultures

During development, individual neurons connect to one another forming neuronal networks that produce distinctive patterns of synchronized activity. Such patterns play a critical role in shaping the network structure and function in the developing brain (38). Upon observing alterations in functional and morphometric properties on the level of individual neurons, we determined if integrated neural circuit activity would be perturbed in RTT cultures. Isogenic WT and RTT *Ngn2* neurons (WIBR3, PGPC14 and CLT-L124W) were co-cultured with mouse astrocytes on 12-well MEA plates with 64 channels in an 8×8 grid per well (Axion Biosystems) (Figure S4A). Every plate was derived from an independent differentiation from *Ngn2*-inducible iPSCs/ESCs and contained 3-6 wells of each isogenic genotype as technical replicates. Time zero was the day when neurons and astrocytes were plated together in co-culture, and five minutes of extracellular recordings were collected each week from week 3 through week 7.

We observed the emergence over time of high frequency neural activity in multiple electrodes similar to those observed in previous experiments (20). Raster plots of action potentials detected using Axion software show synchronous bursts across multiple electrodes for the WIBR3, PGPC14 and CLT isogenic pairs (Figure 4A). As anticipated, we detected variability of firing regimes across the different WT lines, which stresses the importance of using isogenic controls for assessing disease-associated phenotypic changes in human stem cell derived neural networks (39). The number of electrodes with active signals increased over the time course of the recordings likely as a result of development of new connections within the network. We used these metrics to assess the quality and distribution of the neurons in the wells on each plate, as we observed plate-to-plate variability in numbers of active electrodes (Figure S4A,B). Wells that reached a cut-off of at least 32 out of 64 electrodes active over the time course of the recordings were included in analyses to reduce variability. With this cut-off, we could include wells that developed strong network activity and exclude wells that had poor adherence or peeled from the plate. Additionally, at the final time point neurons were treated with the AMPAR antagonist CNXQ to show that spiking activity was abolished (Figure S4C). This supported that synaptic transmission was responsible for network activity patterns.

**Figure 4.**
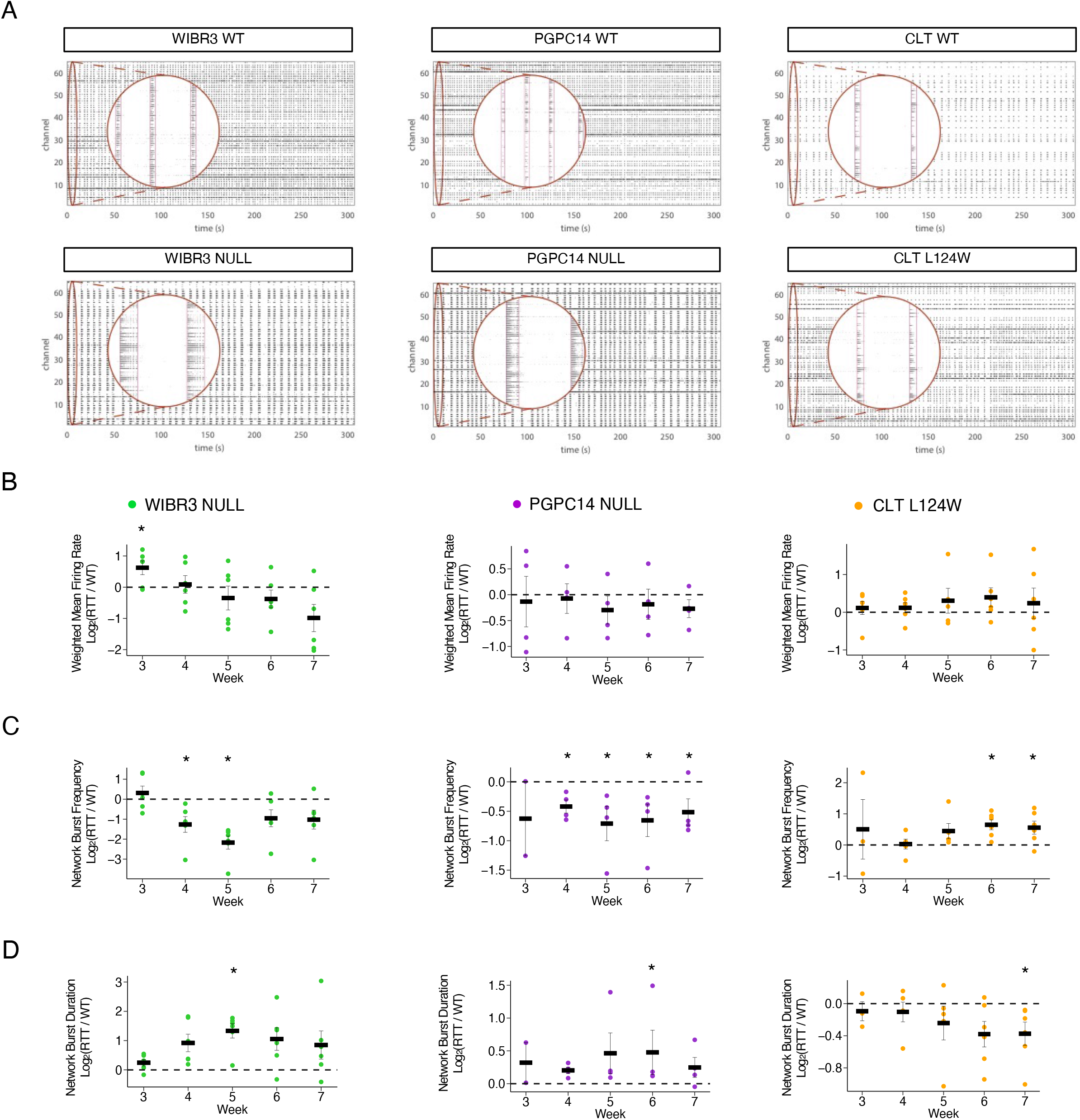
MECP2 null Ngn2 neurons exhibit altered network bursting activity. **A.** Representative 300 second MEA raster plots of spontaneous action potential spikes (black tick marks) from WT and RTT Ngn2 neurons co-cultured on mouse astrocytes at 5 weeks, detected across grid of 64 electrode channels. Inset (red circle) show spikes across multiple channels within synchronous network bursting events. **B.** Quantification of MEA activity shown as log fold change of RTT relative to WT for weighted mean firing rate, **C.** network burst frequency, and **D.** network burst duration. 6 replicate plates of CLT, 6 replicate plates of WIBR3 and 4 replicate plates of PGPC14 isogenic pairs, with 3-6 wells per genotype per plate. All data were shown as mean +/- SEM, * p < 0.05. Statistical significance was evaluated by Student’s t-test or Mann Whitney test, as appropriate, in Figure B-E. *p < 0.05.

Based on the mild electrophysiological and morphological differences in CLT L124W neurons, we predicted a similarly small change in variables that measure network activity. To account for differences in maturation dynamics between plates (20) we normalized the fold-change of RTT values to the respective WT isogenic values within a plate. We examined 6 biological replicate plates of CLT isogenic neurons. The CLT L124W *Ngn2* neurons did not show significant differences in weighted mean firing rate (normalized mean firing rate across active electrodes normalized to the number of active electrodes per well) relative to WT. At late timepoints, increased network burst frequency and decreased network burst duration were observed (Figure 4B-D, right). To detect circuitry defects in *MECP2* null neurons, we then examined 6 biological replicate MEA plates of WIBR3 and 4 biological replicate MEA plates of PGPC14 isogenic pairs. The RTT values are shown as relative log fold change compared to the WT baseline (normalized WT level indicated by dashed horizontal line). The weighted mean firing rate did not have a consistent change in *MECP2* null neurons relative to WT (Figure 4B). However, the *MECP2* null network bursts, showed a consistent decrease in frequency relative to WT beginning at the 4 week time-point. These decreased network burst frequencies were followed by longer network burst durations that transiently emerged at weeks 5-6 in WIBR3 and PGPC14 isogenic lines (Figure 4C,D & by replicate plate in S4D,E). The WIBR3 null isogenic lines had larger fold changes in network bursting properties compared to the PGPC14 null isogenic lines. In contrast, CLT L124W had only late time points with increased network burst frequency or decreased network burst duration that were distinct relative to those shown by *MECP2* null networks.

### Modeling changes in peak bursting frequency in RTT *Ngn2* neural networks

Variations in the frequency of network bursts capture differences between RTT and WT, particularly for the *MECP2* null phenotypes showing more pronounced changes in cellular and network profiles. In order to better characterize such changes in the frequency domain and search for more subtle differences between the different network phenotypes we extracted raw spiking data and performed power spectral density (PSD) estimations on spike trains from each electrode within a well. The PSD calculated directly on recorded spike trains in all 64 channels exhibits clear peaks shown in heatmaps of the frequency of these synchronous network bursts (Figure 5A). When the distributions of peak bursting frequencies were aggregated from weeks 2-7 (Figure 5B and S5A) or normalized to their respective isogenic control (Figure 5C), the WIBR3 and PGP14 nulls were reduced in cumulative median peak bursting frequency across all timepoints. The peak bursting frequency of CLT L124W was slightly increased. Our PSD estimates consistently distinguished *MECP2* null from WT *Ngn2* neural networks, and captured the difference between the direction of peak burst frequency change in the CLT L124W compared to *MECP2* null neurons.

**Figure 5.**
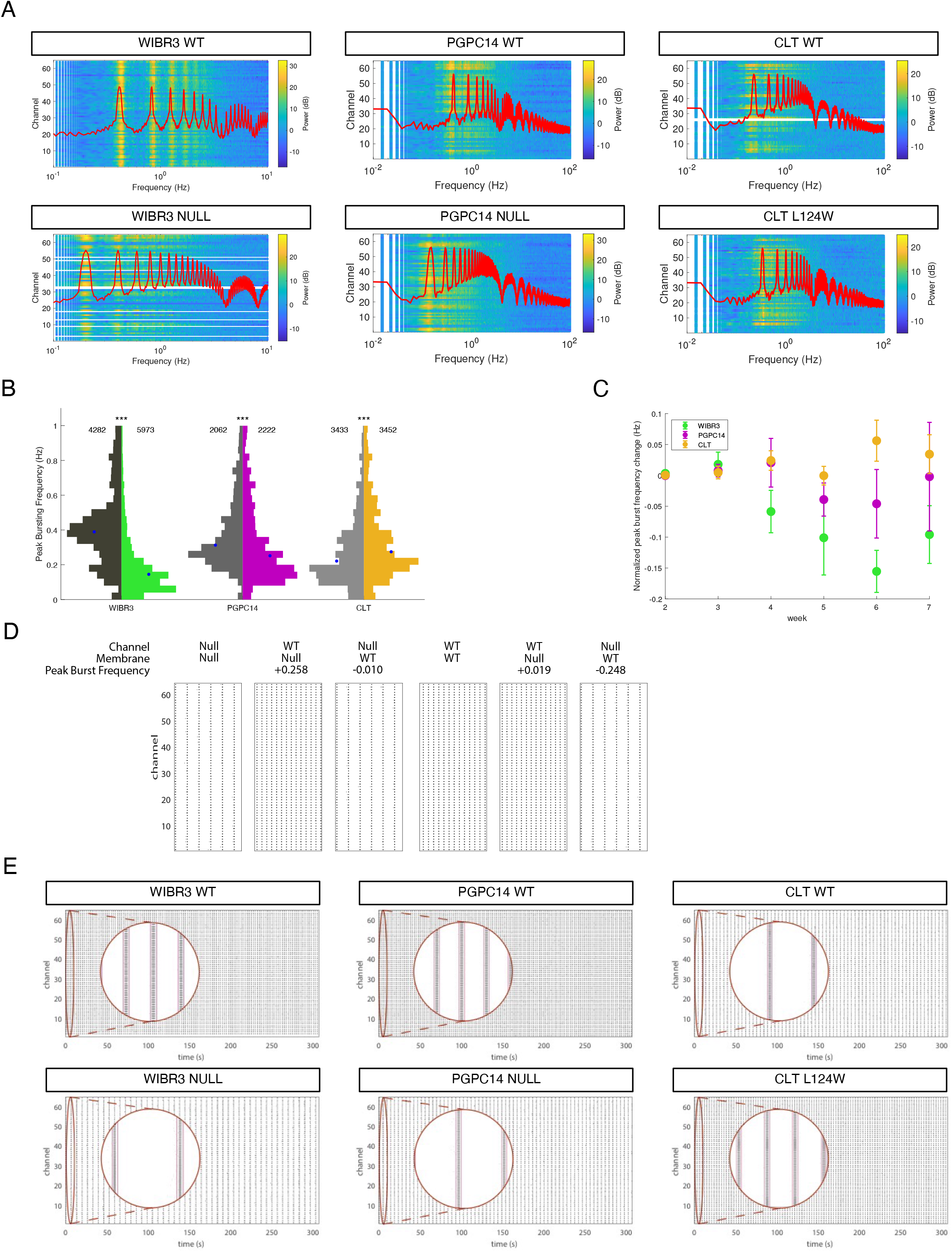
RTT Ngn2 neurons display changes in bursting dynamics. **A.** Peak bursting frequency distribution aggregated from MEA recordings of WT and RTT Ngn2 neurons from week 2-7. 6 replicate plates of WIBR3, 4 replicate plates of PGPC14, and 6 replicate plates of CLT, with 3-6 wells per genotype per plate. Indicated are number of electrodes analyzed, blue dot represents median, *** p < 0.001. **B.** Normalized change in peak frequency from WIBR3 NULL, PGPC14 NULL and CLT 124W Ngn2 neurons across weeks 2-7 shows a developmental trajectory and clear differences between mutant subtypes. **C.** Power spectra of spike trains from recorded WT and RTT *Ngn2* neurons (heatmap), along with the average power spectrum across 1,000 simulated neurons in the spiking network model (red line). Peaks in the experimental data (heatmap) and the computational model (red lines) match well. **D.** Computational drop-out experiment to understand how Na^+^ and K^+^ channel values vs. capacitance, resting membrane potential values contribute to the frequency changes. Change is relative to WIBR Null (with respective Null channel, capacitance, and resting membrane potential values) and WIBR WT (with respective WT channel, capacitance, and resting membrane potential values).Statistical significance was evaluated by Mann Whitney test in Figure A. *p < 0.05, **p < 0.01, ***p < 0.001.

Changes in burst frequency can have at least two different origins, one is changes in the intrinsic properties of individual neurons, and the second is changes in network synaptic connectivity. The observed changes in individual neuron intrinsic properties may suggest the former origin. We hypothesized that the observed differences in peak burst frequency were based on the core phenotype of altered intrinsic currents in the neuronal membrane in RTT networks compared to WT networks. To test this hypothesis, we developed a computational model of excitatory spiking neurons connected into a network that approximates the iPSC-derived cultures. A network of 1000 excitatory adaptive integrate-and-fire (AdIF) neurons (40, 41) was simulated using membrane parameters derived from the patch-clamp recordings, random intra-network connections, connection probability of 0.05 (resulting in an average of 50 synapses per cell), and random Poisson external input (100 synapses spiking at 20 Hz). This spiking model displayed robust network bursts, the features of which are related to single-neuron properties in the network. We then analyzed simulated spike trains in the same way as the experimental data. Overlay of the peak frequency derived from the top fit AdIF simulations (red line in Fig 5A) demonstrated that this spike *adaptation*-driven model recapitulated both the reduced frequency observed in the *MECP2* null neuron recordings and the increased frequency in CLT L124W neurons.

Could systematic differences in network/synaptic connectivity between RTT and WT networks also cause the observed differences in burst frequency? To address this question, we studied the WIBR network model over a range of subthreshold adaptation values (4 to 10 nS) and connection probabilities (0.05 to 0.35). Over this range, network burst frequency increases with the decreased connection probability typically observed in RTT neuron networks (42), while the value of subthreshold adaptation has a smaller effect (Figure S5B). The effect of connection probability on burst frequency is opposite to that observed in the experimental data, indicating that changes in synaptic connection probability are unlikely to explain our observations. This control simulation is thus consistent with our hypothesis that intrinsic adaptation currents are driving the observed changes in burst frequency.

Finally, to determine the contributions of channel currents relative to membrane properties in driving peak burst frequency patterns in RTT, we performed drop out simulations by replacing the WIBR null values with their isogenic WT values. As shown in Fig 5D, simulation with the Null values gives a low frequency of bursts over the representative 30 second duration. When the Null Na^+^ and K^+^ channel values, represented by the adaptation currents, are dropped out and replaced by the WT values, the simulation predicts that burst frequency is rescued. In contrast, when the Null membrane values (capacitance and resting membrane potential) are dropped out and replaced by the WT values, there is no substantial change to the frequency. Reciprocal simulations where the WT values are dropped out and replaced by the null values also predict that it is only the Null channel currents that drive the slow bursting frequency. The simulation model agrees with the experimental circuitry findings for both the *MECP2* null and CLT L124W neurons, and indicates that the *MECP2* null burst patterns are driven by experimentally validated changes in intrinsic currents.

## DISCUSSION

In this study, we defined a wide phenotypic spectrum of *MECP2* mutation-dependent changes in heterochromatin binding, neuron morphology, electrophysiological properties and network connectivity by modeling RTT neurons *in vitro*. We investigated the effects of a novel *MECP2* L124W MBD variant in comparison with three *MECP2* nulls in isogenic induced excitatory neurons. Our findings uncover core functional and morphological phenotypes present in all lines and a broader set of additional impairments exhibited by null neurons (summarized in Table 1). The core phenotypes included increased input resistance, impaired voltage-gated Na^+^ and K^+^ currents, and reduced dendritic complexity. *MECP2* null neurons consistently displayed depolarized resting membrane potential, reduced cell capacitance accompanied by decreased soma area dendrite length and excitatory synapse density, and had disruptions in excitatory synaptic transmission and in neuronal network burst frequency and duration patterns. Simulation modeling replicated the network pattern changes in RTT and predicts that they are driven by channel currents. Thus, the *MECP2* null phenotypic profile encompassed known RTT-associated changes that validated our induced neuron model and revealed novel insights into the development of excitatory synaptic activity and network circuitry.

**Table 1.**
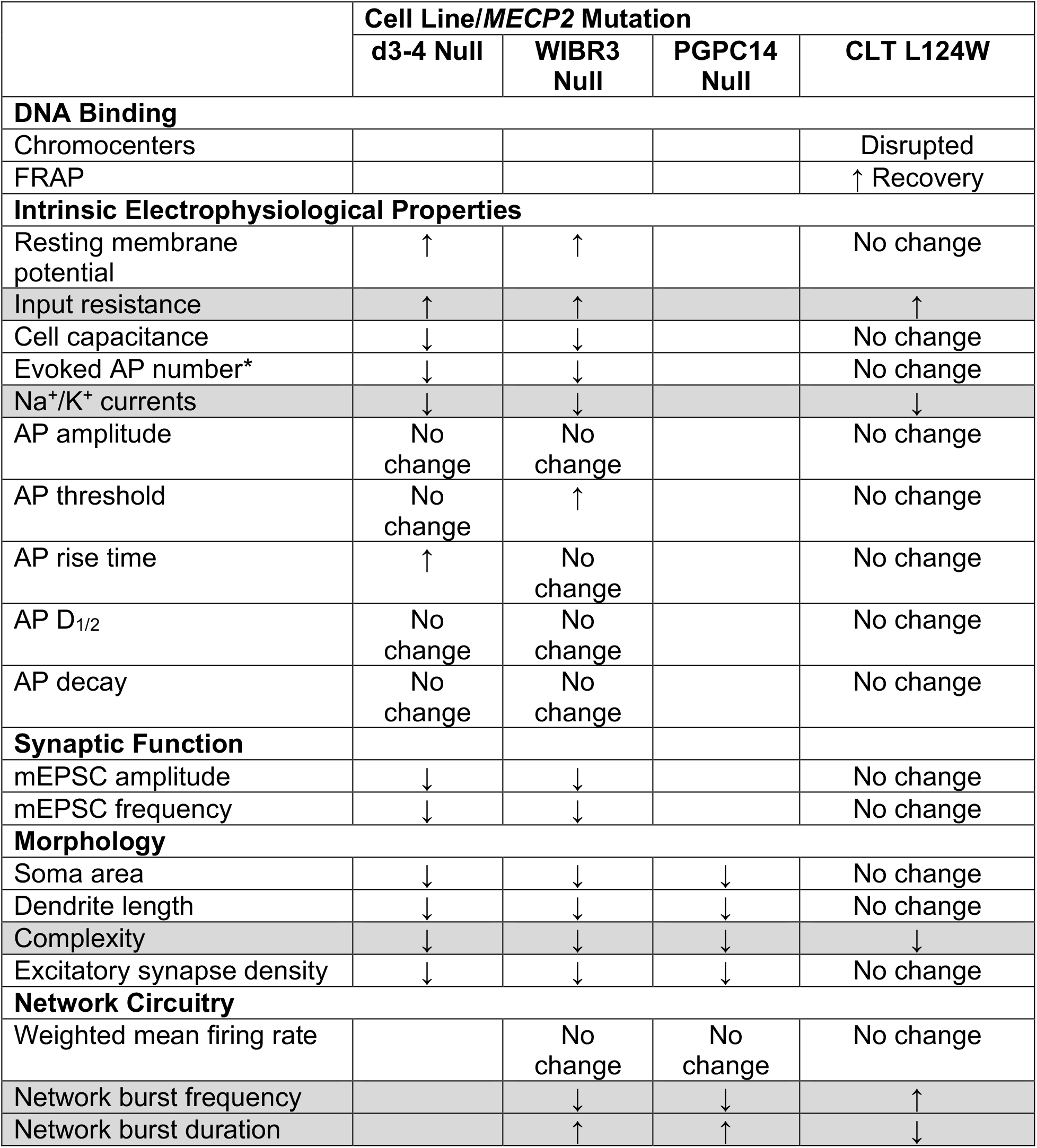
Summary of identified RTT *Ngn2* neuronal phenotypes. Core phenotypes altered in all lines highlighted in gray. *when current injection (over +60 pA).

### L124W cellular and network core phenotypes

The atypical RTT-associated phenotypic profile of L124W MECP2 protein was supported by evidence of partially disrupted chromocenter binding in mouse cells and stable unchanged protein level in induced neurons. One potential experimental artefact we excluded was XCI erosion in the CLT L124W iPSCs. Since expression of the Xi was not reactivated during continued passage, no WT MECP2 was produced to rescue L124W phenotypes. The chromocenter assay was recently used to demonstrate that MECP2 is a component of heterochromatin condensates and that MBD missense mutations compromise this activity consistent with our results (43). Overall, the L124W variant represents a subtype of MECP2 MBD mutation that retains partial function in contrast to the more disrupted heterochromatin found in previous studies of L124F, R106W and other MBD missense variants (7).

We found that CLT L124W neurons showed core phenotype changes in dendrite complexity and intrinsic properties compared to the WT. Importantly, CLT L124W neurons showed increased input resistance. As a consequence of higher input resistance a given depolarizing current will produce a larger change in membrane potential. Thus, increased input resistance explains our finding that the amount of depolarizing current required to evoke an action potential was lower in all mutants as compared with their corresponding controls (Fig. 2E). From this result we conclude that RTT neurons have increased excitability. A recent study on R106W missense iPSC derived neurons also found a graded response in some morphological features and electrophysiological properties compared to MECP2 knockdown neurons, and also did not detect changes in synaptic function that were evident in knockdown neurons (44). In our case we also examined Na^+^ and K^+^ currents in CLT L124W and *MECP2* null neurons, and found these currents are consistently decreased relative to those in WT. As we found limitation of sustained firing of action potentials in response to prolonged current injection in mutant neurons, as compared to WT, we conclude that the decrease in Na^+^ currents predominates over the decrease in K^+^ currents (Fig. 2E). In addition, our network circuitry studies using MEA revealed a core phenotype that network burst frequency and duration are changed, and the direction of this change in CLT L124W and *MECP2* null neurons was successfully modelled computationally. We also recognize the possibility that L124W may have functional effects on other cell types. For example, RTT astrocytes have been shown to exert non-cell autonomous effects on WT and mutant neurons (45, 46). As inhibition was absent in our excitatory neuron model, co-culturing CLT L124W astrocytes with glutamatergic and GABAergic neurons in monolayers or organoids could explore the potential interplay of these signals in single neurons and network activity (47–49).

### *MECP2* null full phenotype and modeling of RTT networks

The *MECP2* null full phenotype included smaller soma size and dendrite length, reduced synapse numbers, and reduced AMPAR-mEPSC amplitude and frequency. Importantly, we observed robust network bursting phenotypes in *MECP2* null neurons. The multi-unit spiking pattern changes revealed that network-wide events occurred less frequently and had extended burst trains in *MECP2* nulls compared to WT. This RTT circuitry phenotype shows similarity to the reduced rate and increased duration of network bursts observed in Kleefstra syndrome induced excitatory neural cultures cocultured with rodent astrocytes, where these changes were found to be mediated by upregulation of N-methyl-D-aspartate receptor subunit *GRIN1* expression (21). Highly perturbed RTT-associated oscillatory network activity phenotype in local field potential MEA recordings of isogenic WT and *MECP2* knockout iPSC-derived cortical organoids showed complete absence of network events (48).

A possible explanation for changes in network dynamics are changes in the intrinsic properties of neurons that may impact their ability to generate bursting patterns. Our spiking network model examined this issue. Model simulations indicate that the changes in RTT null networks bursting frequency may be mediated by neuronal adaptation currents such as inactivation of depolarizing sodium currents, and activity dependent (i.e. calcium dependent) activation of slow hyperpolarizing potassium channels (40, 50), which may be downstream targets of MECP2 (14, 51). This is supported by the decrease in potassium currents seen in RTT neurons (Figure 2). We have also previously found that *MECP2* null neurons derived from NPCs have altered protein levels of K*^+^* channels and AMPA receptors (52). Finally, the computational drop-out simulations support the hypothesis that changes in intrinsic adaptation currents may be the driving variable of the observed changes in burst frequency. Because adaptation currents are known to be critical in shaping neuronal excitability, gain, and responses to sensory input, these changes could play important roles in cortical function (41, 53).

### Significance

In summary, we used a multilevel approach to identify a range of RTT-associated phenotypes relating to form and function in isogenic human stem cell-derived excitatory neurons harbouring a novel L124W missense or *MECP2* null mutations. Our simulations demonstrate the utility of spiking models for analyzing changes in stem cell-derived neuron networks across WT and disease states. The striking network burst frequency changes revealed in *MECP2* null neurons provide a potentially screenable phenotype for testing candidate compounds to rescue the RTT-associated changes in this *in vitro* assay to moderate disease. Full rescue might require correction of protein translation regulation and thus of the channel current deficits found in the core phenotypes.

## METHODS

### Chromocenter and FRAP assays

All imaging and image analysis were performed as previously described (29, 54), with the following exception. FRAP Time-lapse imaging: A series of confocal time lapse images of frames (512 x 512 pixels) imaged at 488nm laser excitation with 0.05 transmission were used to record GFP-tagged protein post bleach recovery. FRAP assay was recorded with a minimum of nine pre-bleach frames, one frame after 1000 µs bleaching with 405 nm laser line at 100% transmission, and 90 post-bleach frames were recorded at equal time intervals.

### Ethics and iPSC reprogramming

iPSCs were reprogrammed under approval of the Canadian Institutes of Health Research Stem Cell Oversight Committee and the Research Ethics Board of The Hospital for Sick Children (REB #1000050639) and the University of California San Diego (IRB #14123ZF). Existing iPSC lines are listed in Table S1.

CLT iPSC lines were generated from fibroblasts. Fibroblasts were collected from a small incision in the skin after a skin biopsy and tissue was washed in PBS 1x with 4% penicillin/streptomycin (Gibco). Later, the tissue was broken into small pieces using needles and plated in culture media (DMEM/F12, 15% FBS, 1% MEM NEAA, 1% L-glutamine and 1% penicillin/streptomycin). After 1 month, fibroblasts migrated out from the tissue. After reaching high confluency, cells were expanded or frozen as stock (55, 56). Fibroblasts were reprogrammed using Sendai virus (Invitrogen) in feeder-free conditions on Matrigel (Corning) with mTeSR1 (STEMCELL Technologies). After colonies started to grow, colonies were picked manually and expanded on feeder-free Matrigel-coated dishes and frozen with CryoStore (STEMCELL Technologies). Mycoplasma contamination was routinely checked.

### iPSC/ESC culturing and maintenance

Human iPSC/ESCs (Table S1) were maintained on dishes coated with Matrigel in mTeSR1 or StemMACS iPS-Brew XF (Miltenyi Biotec) media with 1% penicillin/streptomycin. iPSCs were passaged weekly with ReLeSR (STEMCELL Technologies) with daily media changes except for the day following passaging. Accutase (Innovative Cell Technologies) and media with 10 μM Rho-associated kinase (ROCK) inhibitor (Y-27632, STEMCELL Technologies) was used for any single-cell dissociation. Routine mycoplasma testing was performed.

### Karyotyping

iPSC karyotyping and standard G-banding chromosome analysis (400-banding resolution) was performed by The Centre for Applied Genomics (TCAG) at The Hospital for Sick Children.

### Pluripotency assays

iPSC colonies were fixed and stained with pluripotency markers OCT4, NANOG, SSEA4 and TRA-1-60. Following embryoid body formation and spontaneous 3-germ layer differentiation for 16 days, resulting cells were fixed and stained for α-SMA, AFP and β-III-tubulin. To assess gene expression profile of iPSCs, RNA was extracted from 1 confluent well of a 6-well plate using RNA PureLink RNA mini kit (Life Technologies) as per manufacturer’s protocol. Samples were sent to TCAG at the Hospital for Sick Children for Bioanalyzer assessment and sequenced as paired end 2×125 bases on an Illumina HiSeq 2500 at a depth of 15M reads. Resulting FASTQ files were uploaded to www.pluritest.org for analysis (34). Files can be found with GEO accession GSE148435.

### XCI assays

Androgen receptor (AR) assay was performed as previously described (9). In brief, 400 ng of genomic DNA from iPSCs/ESCs was digested by CpG methylation-sensitive restriction enzymes *Hpa*II and *Hha*I for 6 hours. 2 μL of digestion was amplified using Platinum Taq DNA Polymerase High Fidelity (Invitrogen) with AR gene primers with a FAM label (Table S2). Male samples were used as a control to confirm complete digestion. Electrophoresis was performed by TCAG at The Hospital for Sick Children. Traces were analyzed using PeakScanner (Thermo Fisher Scientific). To identify which *MECP2* allele was expressed on the active X chromosome, RNA was isolated from iPSCs and reverse transcribed into cDNA using SuperScript III (Invitrogen). Primers flanking the variant region (Table S2) were used for amplification with Platinum Taq DNA Polymerase High Fidelity or Q5 High Fidelity DNA Polymerase (New England BioLabs), followed by cloning as per the TOPO TA Cloning Kit (Invitrogen) or Zero Blunt TOPO Cloning Kit (Invitrogen) in OneShot TOP10 (Thermo Fisher Scientific) or Max Efficiency DH5α (Invitrogen) competent *E. coli* strains. 5-10 bacterial colonies per iPSC cell line tested were picked and grown in LB Medium (MP Biomedicals) for DNA extraction with Quick Plasmid MiniPrep Kit (Invitrogen). Samples were Sanger sequenced at TCAG and aligned to WT *MECP2* template using benchling.com.

### CRISPR/Cas9 gene editing

The *MECP2* indel mutation was generated using a previously described gene editing strategy (32). Briefly, pSpCas9(BB)-2A-Puro (PX459) V2.0 (Addgene #62988) was cloned with an oligonucleotide designed to target *MECP2* exon 3 using benchling.com CRISPR prediction tool (described in Table S2). Indels were introduced by transfecting 8×10^5^ cells with 1.5 μg plasmid in 100 μl scale using the Neon Transfection System (ThermoFisher) with one pulse at 1500 millivolts for 30 milliseconds (∼40-70% transfection efficiency). Transfected cells were plated in 2 wells of a 6-well plate with mTeSR1 media supplemented with CloneR (STEMCELL Technologies) to enhance cell survival. From D2-5, puromycin (0.5 ug/ml) was added to daily to mTeSR1 media changes and surviving single colonies were grown to D12-18 before transferring to 24-well plates. Isolated clones were passaged and gDNA was harvested with a Quick DNA miniprep kit (Zymo) and PCR products were sent for Sanger sequencing by TCAG at the Hospital for Sick Children.

### Lentivirus generation and transduction

*FUW-TetO-Ngn2-P2A-EGFP-T2A-puromycin* and *FUW-rtTA* plasmids for excitatory cortical neuron differentiations were kindly gifted by T. Sudhof (36) and packaged as per a previously described protocol (57). Lentiviruses were produced in HEK293T cells using a third generation packaging system (*pMDG.2*, *pRSV-Rev* and *pMDLg/pRRE*). Viral particles in the supernatant were collected 48 hours post-transfection, filtered and concentrated by 91,000 x g centrifugation for 2 hours at 4°C. For iPSC transductions, accutase (Innovative Cell Technologies) was used to obtain 5×10^5^ cells single-cells per well of a Matrigel-coated 6-well plate that were incubated with concentrated virus in mTeSR1 with 10 μM ROCK inhibitor and 8 μg/ml polybrene (Sigma) for 8 hours. Media was changed to mTeSR1 and transduced iPSCs were maintained and expanded to acquire frozen stocks.

### *Ngn2* neuronal direct conversion

Excitatory cortical neurons were generated as previously described (32, 35). Briefly, on Day 0 iPSCs were made into single-cells by accutase and plated at 3×10^5^-1×10^6^ cells per well on Matrigel-coated 6-well plates with StemMACS iPS-Brew XF media supplemented with 10 μM ROCK inhibitor. All media mentioned for Day 1-8 contained 1x penicillin/streptomycin (Gibco), 1 μg/mL laminin (Sigma), 2 μg/mL doxycycline hyclate (Sigma), 10 ng/ml BDNF (Peprotech) and 10 ng/mL GDNF (Peprotech). On Day 1, media was changed to CM1 consisting of DMEM/F12 (Gibco), 1x N2 (Gibco) and 1x NEAA (Gibco), supplemented with 10 μM ROCK inhibitor. On Day 2, media was changed to CM1 supplemented with 2-5 μg/mL puromycin (Sigma). On Day 3, media was changed to CM2 consisting of Neurobasal (Gibco), 1x B27 (Gibco) and 1x Glutamax (Gibco), supplemented with 2-5 μg/mL puromycin. On Day 4-5, media was changed to CM2. On Day 6, media was changed to CM2 supplemented with 10 μM Ara-C (Sigma). On Day 8, doxycycline induction was withdrawn and neurons were re-seeded into new dishes in CM2 and co-cultured with P1 mouse astrocytes unless otherwise noted for downstream assays.

### Protein quantification

Protein was extracted from neurons (not grown with mouse astrocytes) washed in ice-cold PBS using radioimmune precipitation assay (RIPA) buffer (25 mM Tris-HCl, pH 7.6, 150 mM NaCl, 1% Nonidet P-40, 1% sodium deoxycholate and 0.1% SDS). Equivalent protein masses were loaded into capillaries for detection using the 12-230 kDa Separation Kit for the Wes (ProteinSimple). Multiplexed probing of MECP2 and β-actin by chemiluminescence was quantified using Compass for SW software (ProteinSimple).

### Patch-clamp electrophysiology

Electrophysiological recordings were made as reported previously (10, 37). Briefly, whole-cell patch-clamp recordings were performed with an Axopatch 1-D amplifier (Molecular Devices, USA) operated with Clampex 9.2 software and a DigiData 1200 series interface (Molecular Devices, USA), and the currents were sampled at 10 kHz and filtered at 2 kHz. The recordings were analyzed off-line using Clampfit 9.2 software (Molecular Devices, USA). All electrophysiological experiments and data analysis were conducted with *Ngn2* neuron samples blinded to the investigator. To examine intrinsic membrane properties, the recordings were performed in a 35-mm tissue culture dish filled with an extracellular solution containing (in mM): 140 NaCl, 5.4 KCl, 1 MgCl_2_, 15 HEPES, 2 CaCl_2_, 10 glucose, and the solution was adjusted to pH 7.35 with NaOH. Patch-clamp pipettes (5 to 8 MΩ) were pulled from capillary glass (1.5 mm diameter; World Precision Instruments, USA) using a P-87 pipette puller (Sutter Instrument Co., USA), and filled with an intracellular solution compose of (in mM): 144 K^+^-gluconate, 10 KCl, 2 Mg-ATP, 2 EGTA, 10 HEPES. The solution was adjusted to pH 7.2 with KOH. Voltage-gated ion currents, under the whole-cell voltage-clamp condition, were elicited at the holding membrane potential of −70 mV by stepping the membrane potential to a series of potentials from –80 mV to +60 mV for 400 ms. Action potentials, under current-clamp condition, were evoked from the membrane potential of around −75 mV by injecting a series of current steps from −5 pA to 100 pA (in 5 pA increments). Data from some neurons in both WT and mutant lines, with 10 pA increments, or maximum current injection of less than 100 pA, were pooled with those obtained with the current injection protocol above since there was no difference. In current-clamp, we subtracted a liquid junction potential of 16 mV from the membrane potential values measured with K^+^-gluconate based intracellular solutions.

To investigate synaptic function, mEPSCs, in voltage-clamp, were recorded human *Ngn2* neurons at the holding membrane potential of −60 mV. Recording micropipettes were filled with an intracellular solution as described above. The extracellular recording solutions consisted of (in mM): 140 NaCl, 1 MgCl_2_, 5.4 KCl, 1.3 CaCl_2_, 25 glucose, 15 HEPES, 0.003 glycine, 0.001 strychnine, 0.01 bicuculline, 0.0005 tetrodotoxin, and pH was adjusted to 7.35 with NaOH. mEPSCs were detected and analyzed off-line using Mini Analysis Program (Synaptosoft Inc, USA).

### Neuron sparse labeling

Neurons were transfected with *pL-SIN-EF1α-eGFP* plasmid (Addgene 21320) using 1 µg of DNA and 2 µL of Lipofectamine 2000 (Thermo Fisher Scientific) per well of a 24-well plate. 48 hours post-transfection, neurons were fixed at 6 weeks and stained for MAP2 and GFP.

### Immunocytochemistry and imaging

iPSCs were fixed with 4% formaldehyde in PBS for 8 minutes at room temperature. Neurons grown on 12mm round glass coverslips (Bellco) or 24-well µ-Plates (Ibidi) were fixed with 4% formaldehyde in 0.4 M sucrose Krebs buffer for 8 minutes to preserve cytoskeletal integrity (58). Fixed cells were permeabilized with 0.1% Triton X-100 for 8 minutes at room temperature and blocked for at least 1 hour at room temperature in ICC buffer (10% normal goat (Cedarlane) or donkey (Sigma) serum in PBS with 0.05% Tween 20). Primary antibodies were diluted in ICC buffer and incubated overnight at 4°C. Secondary antibodies were diluted in PBS with 0.05% Tween 20 and incubated for 1 hour at room temperature. Unless otherwise indicated, nuclei were counterstained with 1 µg/ml DAPI in PBS for 5 minutes at room temperature. If cells were on coverslips, they were rinsed in molecular grade water and mounted on slides in Prolong Gold antifade mounting media (Thermo Fisher). Details for primary antibodies used in this study can be found in Table S3.

iPSC pluripotency and 3-germ layer assays were imaged with a Leica DM14000B epifluorescence microscope using a DFC7000T camera and LAS X software. Neurons for soma/dendrite tracing were imaged with an Olympus 1X81 spinning disk confocal microscope using a Hamamatsu C9100-13 EM-CDD camera and Volocity software (Perkin Elmer). Synapses were imaged with a Leica SP8 Lightning Confocal/Light Sheet point scanning confocal microscope and LAS X software.

### Image analysis

Blinded images were analyzed using Filament Tracer in Imaris (Bitplane) by two individuals. Soma were manually traced with semi-automated reconstruction of dendrites on sparsely GFP labeled neurons that were co-stained with MAP2 (3-5 replicate batches, total = 404 neurons). Synapse density was manually counted for over-lapping SYN1 and HOMER1 puncta along a given length of MAP2 dendrite signal (2 replicate batches, total = 240 dendrite segments).

### MEA plating and recording

12-well CytoView MEA plates (Axion Biosystems) with 64 channels per well in 8×8 electrode grids were coated with filter sterilized 0.1% poly(ethylenimine) solution (Sigma) in borate buffer pH 8.4 for 1 hour at room temperature, washed 4 times with water and dried overnight. Day 8 differentiated *Ngn2* neurons were re-seeded at a density of 1×105 cells per 100 µL droplet in CM2 media consisting of BrainPhys (STEMCELL Technologies), 1x penicillin/streptomycin, 10 ng/ml BDNF and 10 ng/mL GDNF, supplemented with 400 μg/mL laminin and 10 μM ROCK inhibitor. Droplet of cells was applied directly on top of electrodes in each well, incubated for 1 hour at 37°C with hydration and then followed by slow addition of CM2 media supplemented with 10 μg/mL laminin. 24 hours later, 2×104 P1 mouse astrocytes were seeded on top of the neurons in each well. CM2 media supplemented with 10 μg/mL laminin was changed consistently 24 hours prior to recording twice per week from 2 weeks onwards on the Maestro MEA platform (Axion Biosystems). Each plate was incubated for 5 minutes on a 37°C heated Maestro device, then recorded for 5 minutes of spontaneous neural activity using AxIS 2.0 software (Axion Biosystems) with a 0.2-3 kHz bandpass filter at 12.5 kHz sampling frequency. Spikes were detected at 6x the standard deviation of the noise on electrodes. Offline analysis used the Neural Metric Tool (Axion Biosystems), where an electrode was considered active if at least 5 spikes/min was detected. Poisson surprise burst and Envelope network burst algorithms with a threshold of at least 25% active electrode were applied. Further analyses and normalization of MEA metrics were completed in RStudio and MATLAB.

### Statistical analyses

Statistical tests for assays were performed in RStudio (v1.1.423), SigmaPlot, GraphPad Prism, or MATLAB. D’Agostino & Pearson test was used to determine normality for datasets. Comparisons within respective isogenic pairs were conducted by two-way repeated measure ANOVA, Student’s t, one sample t or Mann-Whitney tests where appropriate (* p < 0.05, ** p < 0.01, *** p < 0.001).

### Network model

The network model consists of N adaptive leaky integrate-and-fire (ADIF) neurons. Consistent with the biological iPSC-derived neuronal cultures studied here, the network is fully excitatory. The membrane potential *V_i_* of the th unit evolves in time according to the equations:

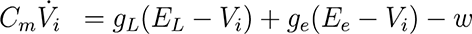

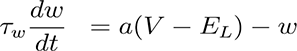

where *C_m_* is the membrane capacitance, *g_L_* is the leak conductance, *E_L_* is the resting membrane potential, *g_e_* is the time-dependent excitatory synaptic conductance, and *E_e_* is the excitatory reversal potential. In the second equation, *ω* is the adaptation variable, *τ_ω_* is the adaptation time constant, and *α* controls subthreshold adaptation.

When the membrane potential of neuron *i* exceeds threshold *V_th_*, the neuron emits a spike, and the following reset conditions occur:

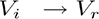

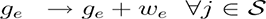

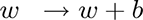

where *V_r_* is the reset potential, *ω_e_* is the excitatory synaptic weight, *S* is the set of neurons postsynaptic to neuron *i*, and *b* is the adaptation step current. Following the initial conductance increase caused by an incoming spike, the excitatory and inhibitory synaptic conductances follow simple exponential decay:

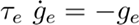

where *τ_e_* is the excitatory synaptic time constant. Populations of *N* = 1000 neurons were simulated for 300 seconds, representative of the *in vitro* culture recordings, with time-step of 0.1 ms in the Brian2 simulator (59). Neurons were randomly connected with excitatory synapses and a connection probability of 0.05, leading to an average of 50 synapses per cell in the simulated networks.

### Burst Frequency

MEA raw voltage signals were measured for 300 seconds at a 12.5kHz sampling frequency and Butterworth bandpass filtered between 200Hz and 3000Hz. Spike times were subsequently detected using an adaptive threshold crossing (threshold set at 6 x Std dev) and organized into spike trains per electrode, then further downsampled to 1kHz. The power spectral density (PSD) of each spike train was estimated using MATLAB’s pwelch() function, and was expressed in decibels (dB) as a ratio against the PSD of a randomly permuted version of the same spike train. The power spectrum provides a numerical weight for each measurable frequency of an input time series. The described synchronous, regular bursting corresponded to the global peaks of the spectra between 0 Hz and 1 Hz. Spectra with global peaks outside this range were found to not clearly exhibit such bursting and were therefore excluded from analysis.

### Network Scan

To identify the ideal time constant for adaptation (*τ_ω_*), we scanned a two-dimensional parameter space of capacitances (*C_m_*) and neuronal membrane time constants (*τ_m_*) in the range observed in patch-clamp recordings. Peak burst frequency in the simulations was evaluated through the same approach as the recorded culture data. Top simulations for the null and atypical CLT L124W conditions were determined as having the smallest absolute difference from the mean peak burst frequency of the recorded data.

To deconstruct the influence of various electrophysiologically-contributing variables to the changes in burst frequency, we performed a computational drop-out experiment. Here we changed either Na^+^/K^+^ currents or capacitance and resting membrane values, based on the reported values of the isogenic pair in Table 4, in the WIBR wildtype and WIBR null networks and calculated peak burst frequency as previous mentioned.

## Data and Code accession

RNAseq datasets are available at GEO accession GSE148435. Code for visualizing electrophysiology, morphology and MEA analysis will be available on GitHub. Code for simulation modeling will be available.

## Reagent Lists

**Table S1.** Description of iPSC/ESC lines.

| Individual | Cell Type | Genotype | # of Clones | *MECP2*<sup>+/-</sup> Mutation Description | Reference |
| --- | --- | --- | --- | --- | --- |
| CLT | iPSC | WT | 2 | Wild-type | This study |
|  |  | L124W | 2 | Patient c.371T>G; L124W missense mutation in methyl binding domain |  |
| d3-4 | iPSC | WT | 1 | Wild-type | Cheung et al, 2011 |
|  |  | NULL | 1 | Patient indel disrupting exons 3 & 4 |  |
| WIBR3 | ESC | WT | 1 | Wild-type | Li et al, 2013 |
|  |  | NULL | 1 | TALEN-edited deletion of exons 3 & 4, Clone #913 from Yun Li. |  |
| PGPC14 | iPSC | WT | 1 | Wild-type | Hildebrandt et al, 2019 |
|  |  | NULL | 1 | CRISPR/Cas9-edited 1 bp insertion (frameshift, premature stop codon) | This study |

**Table S2.**
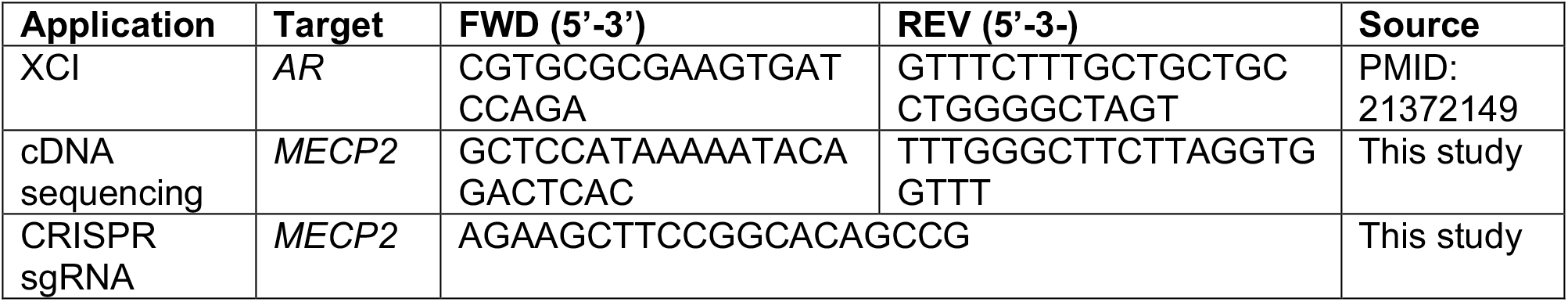
Oligonucleotide sequences.

**Table S3.** Primary antibodies.

| Antibody | Species | Dilution |  | Source | Cat # |
| --- | --- | --- | --- | --- | --- |
|  |  | ICC | Wes |  |  |
| Anti-α-SMA | Mouse | 1:200 | | Invitrogen | 18-0106 |
| Anti-β-actin | Mouse | | 1:500 | Sigma | A5441 |
| Anti-β-III tubulin | Mouse | 1:200 | | Chemicon | MAB1637 |
| Anti-AFP | Mouse | 1:200 |  | R&D | MAB1368 |
| Anti-GFP | Chicken | 1:1000 |  | Thermo Fisher | A10262 |
| Anti-HOMER1 | Guinea pig | 1:500 |  | Synaptic Systems | 160004 |
| Anti-MAP2 | Mouse | 1:1000 |  | Sigma | M1406 |
| Anti-MAP2 | Guinea pig | 1:1000 |  | Synaptic Systems | 188004 |
| Anti-MECP2 | Rabbit | 1:1000 | 1:50 | Cell Signalling | 3456 (D4F3) XP |
| Anti-Nanog | Rabbit | 1:200 |  | Cell Signalling | 4903P |
| Anti-OCT4 | Rabbit | 1:200 |  | Abcam | ab19857 |
| Anti-SSEA4 | Mouse | 1:100 |  | Invitrogen | 41-4000 |
| Anti-SYN1 | Rabbit | 1:200 |  | Millipore | AB1543P |
| Anti-TRA-1-60 | Mouse | 1:100 |  | Invitrogen | 41-1000 |

## ACKNOWLEDGEMENTS

We thank the family of the atypical patient for their participation, Lyndon Duong, Milad Khaki and Fraser McCready for preliminary MEA computational studies; Janice Hicks for the great technical support; Yun Li for the kind gift of WIBR3 WT and NULL cells; TCAG for karyotyping and sequencing; and the SickKids Imaging Facility. This research was funded by grants from Col Harland Sanders Rett Syndrome Research Fund at the University of Toronto (to J.E.), SFARI (Research grant #514918 to J.E. and J.M-T), CIHR (MOP-133423 to J.E. and M.W.S.; ERARE Team Grant ERT161303 to J.E.), CIHR foundation grant (FDN-154336 to M.W.S), Ontario Brain Institute (POND Network to J.E.), McLaughlin Centre Accelerator grant (to J.E.), John Evans Leadership Fund & Ontario Research Fund (to J.E), Beta Sigma Phi International Endowment Fund (to J.E.), BrainsCAN at Western University through the Canada First Research Excellence Fund (CFREF) (to GB, KP, LM, JMT), National Institutes of Health (NIH) grants # R01MH108528, R01MH109885, and R01MH1000175 to ARM, and from Ontario Rett Syndrome Association to JBV. Trainee support was provided by Restracomp (to LCD).

## AUTHOR CONTRIBUTIONS

RSM, WZ, TIS, IRF, ARM, JBV, MWS and JE conceptualized and designed experiments. RSM, WZ, TIS, IRF, LCD, MRH, DCR, MM, WW, AP and JL performed and analyzed experiments. KP, RSM, GB, LM and JM-T conceptualized, designed, performed and analyzed simulations. ARM, LM, JBV, JM-T, MWS and JE supervised and obtained funding. RSM, WZ, TIS, KP, IRF, GB, JM-T, LM and JE wrote the paper. All authors revised and approved the paper.

## Competing interests

A.R.M. is a co-founder and has equity interest in TISMOO, a company dedicated to genetic analysis and brain organoid modeling focusing on therapeutic applications customized for autism spectrum disorder and other neurological disorders with genetic origins. The terms of this arrangement have been reviewed and approved by the University of California San Diego in accordance with its conflict of interest policies. M.R.H. became an employee of STEMCELL Technologies Inc during the preparation of this manuscript.

## SUPPLEMENTAL FIGURE CAPTIONS

**Figure S1.**
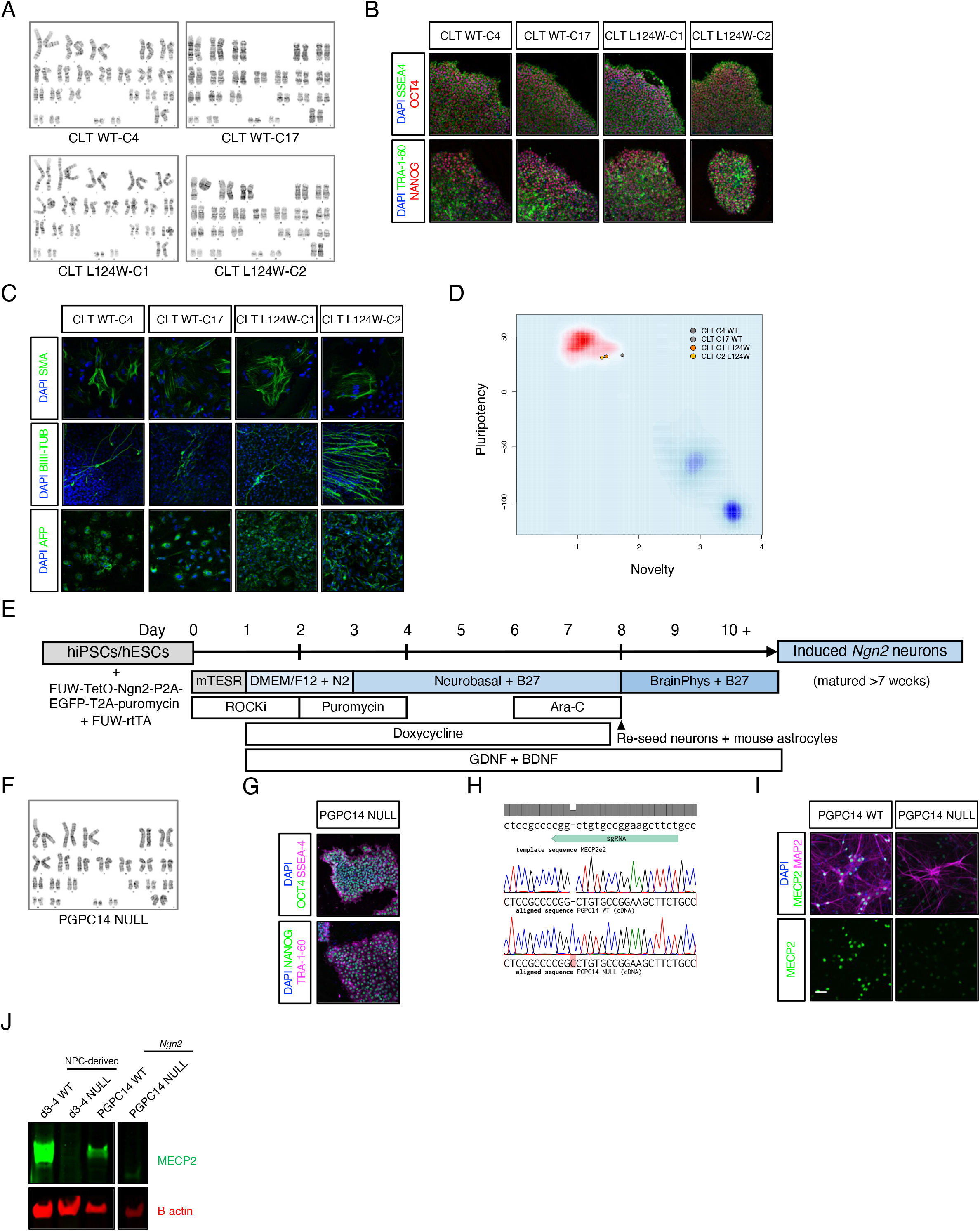
Generation of MECP2 L124W and PGPC14 MECP2 null iPSCs and Ngn2 excitatory cortical neurons. **A.** G-band karyotyping of two CLT WT (C4 and C17) and two CLT L124W (C1 and C2) female iPSC lines (46XX). **B.** Representative images of iPSC colonies from two CLT WT (C4 and C17) and two CLT L124W (C1 and C2) lines stained with pluripotency-associated nuclear markers OCT4 and NANOG (red) and surface markers SSEA4 and TRA-1-60 (green). **C.** Representative images of spontaneous 3-germ layer embryoid body differentiations from two CLT WT (C4 and C17) and two CLT L124W (C1 and C2) iPSC lines stained with mesoderm marker SMA, ectoderm marker β-III-tubulin and endoderm marker AFP (green). **D.** Pluritest clustering of two CLT WT (C4 and C17) and two CLT L124W (C1 and C2) iPSC colonies based on gene expression profile comparison to established iPSC lines. **E.** Schematic of induced *Ngn2* neuronal differentiation protocol from iPSCs/ESCs. **F.** G-band karyotyping of PGPC14 null iPSC line (46XX). **G.** Representative images of iPSC colonies from PGPC14 null line stained with pluripotency-associated nuclear markers OCT4 and NANOG (green) and surface markers SSEA4 and TRA-1-60 (magenta). **H.** Sanger sequencing of cDNA for the 1 bp insertion frameshift mutation (p.Val74CysfsTer16) from restricted expression of the active X chromosome CRISPR/Cas9 gene edited PGPC14 iPSCs. **I.** Representative images of 6 week old PGPC14 WT and null *Ngn2* neurons co-cultured on mouse astrocytes stained with MECP2 (green) and MAP2 (magenta). Scale bar = 50 μm. **J.** Western blot of MECP2 protein in 6 week old PGPC14 WT and null *Ngn2* neurons, compared to 4 week old d3-4 WT and null NPC-derived neurons.

**Figure S2.**
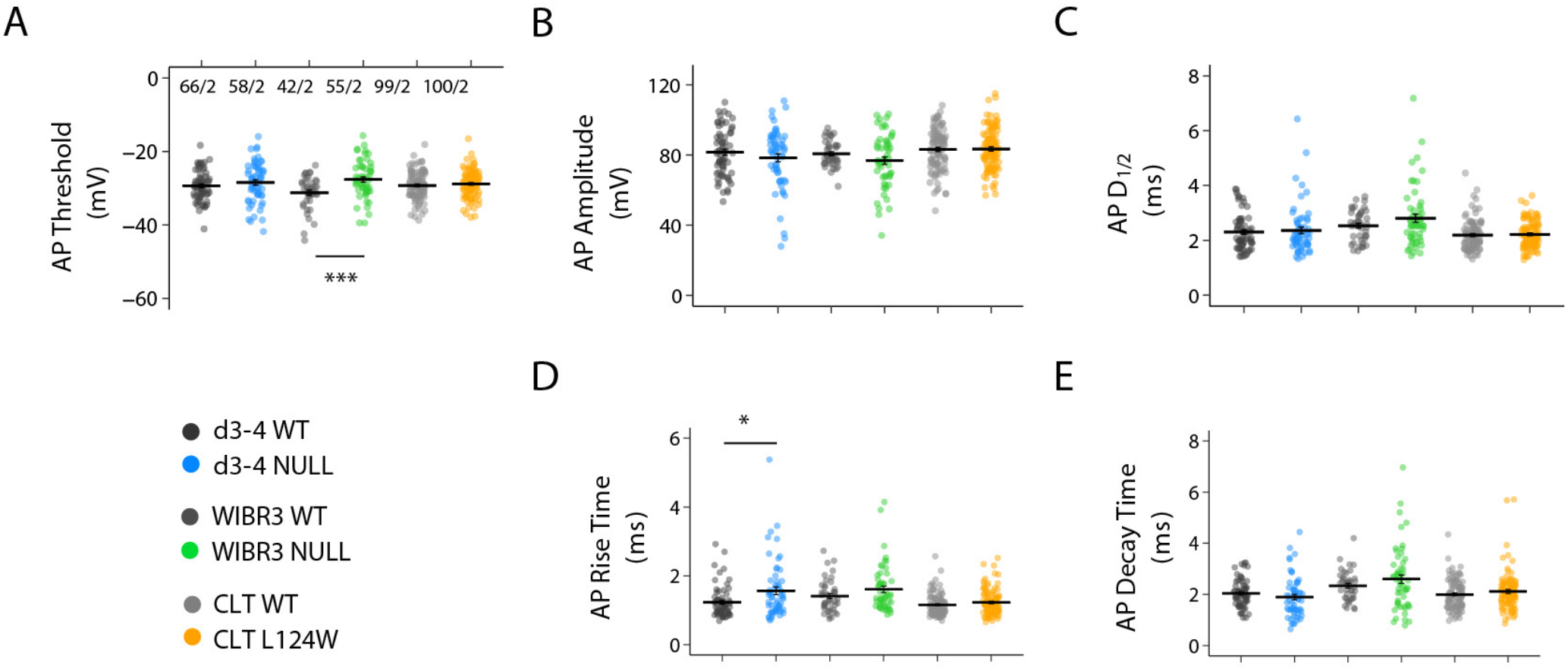
Alterations in the parameters of action potentials in RTT Ngn2 neurons. **A-E.** Scatter plots showing all data points for the threshold, amplitude, rise time, half-duration and decay time of evoked action potentials in 4-5 week-old Ngn2 neurons of WT (n = 66, 42, and 99, respectively) and RTT from d3-4 (n =58), WIBR (n = 55), CLT (n = 100). The graphs also display average parameters of action potentials. The data are shown as mean +/- SEM. Liquid junction potential was not corrected for the action potential threshold as plotted. Statistical significance was evaluated by Student’s *t*-test or Mann Whitney test, as appropriate. *p < 0.05, ***p < 0.001.

**Figure S3.**
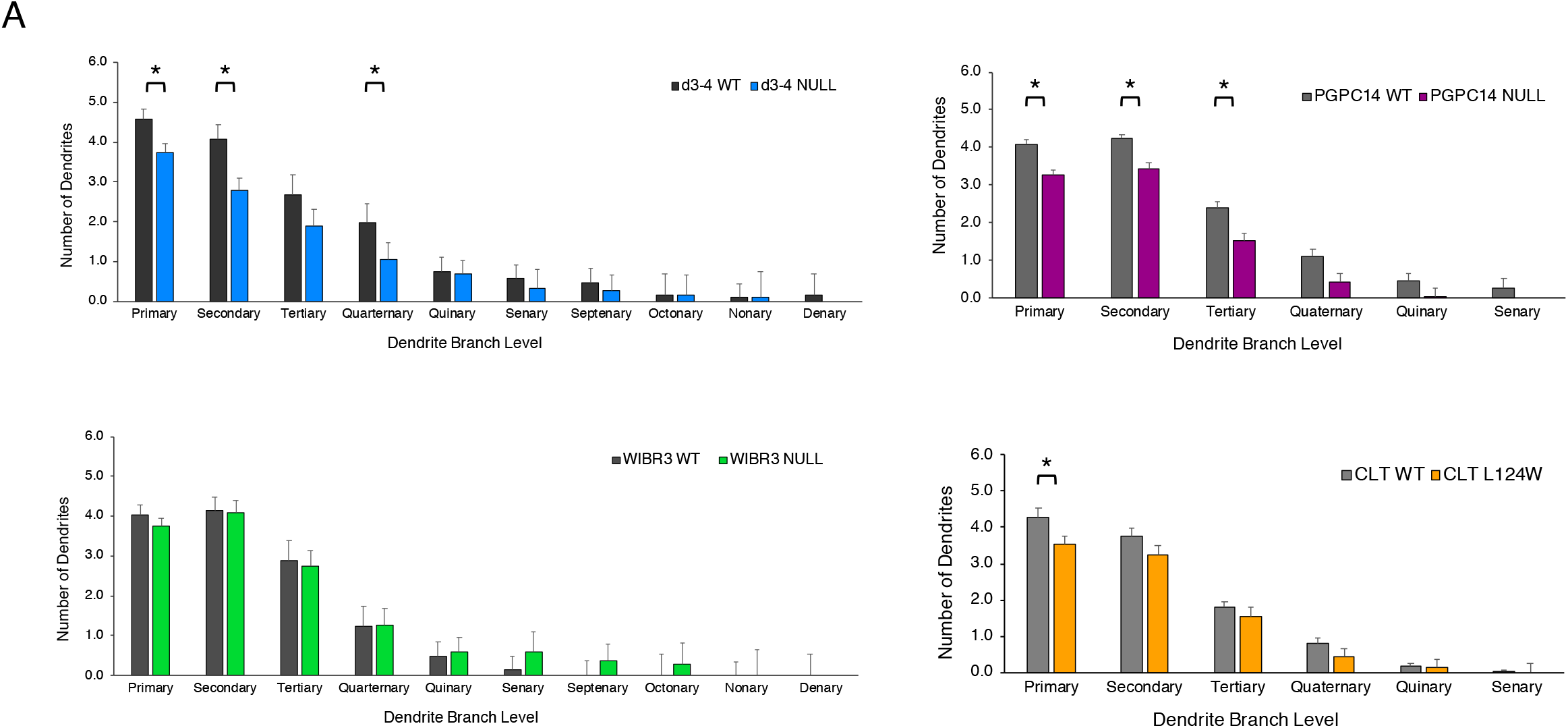
Branching alterations in RTT Ngn2 neurons. **A.** Quantification of 6 week old WT and RTT Ngn2 neurons co-cultured with mouse astrocytes, for dendrite branching order (manual counting). Mean +/- SEM, * p < 0.05.

**Figure S4.**
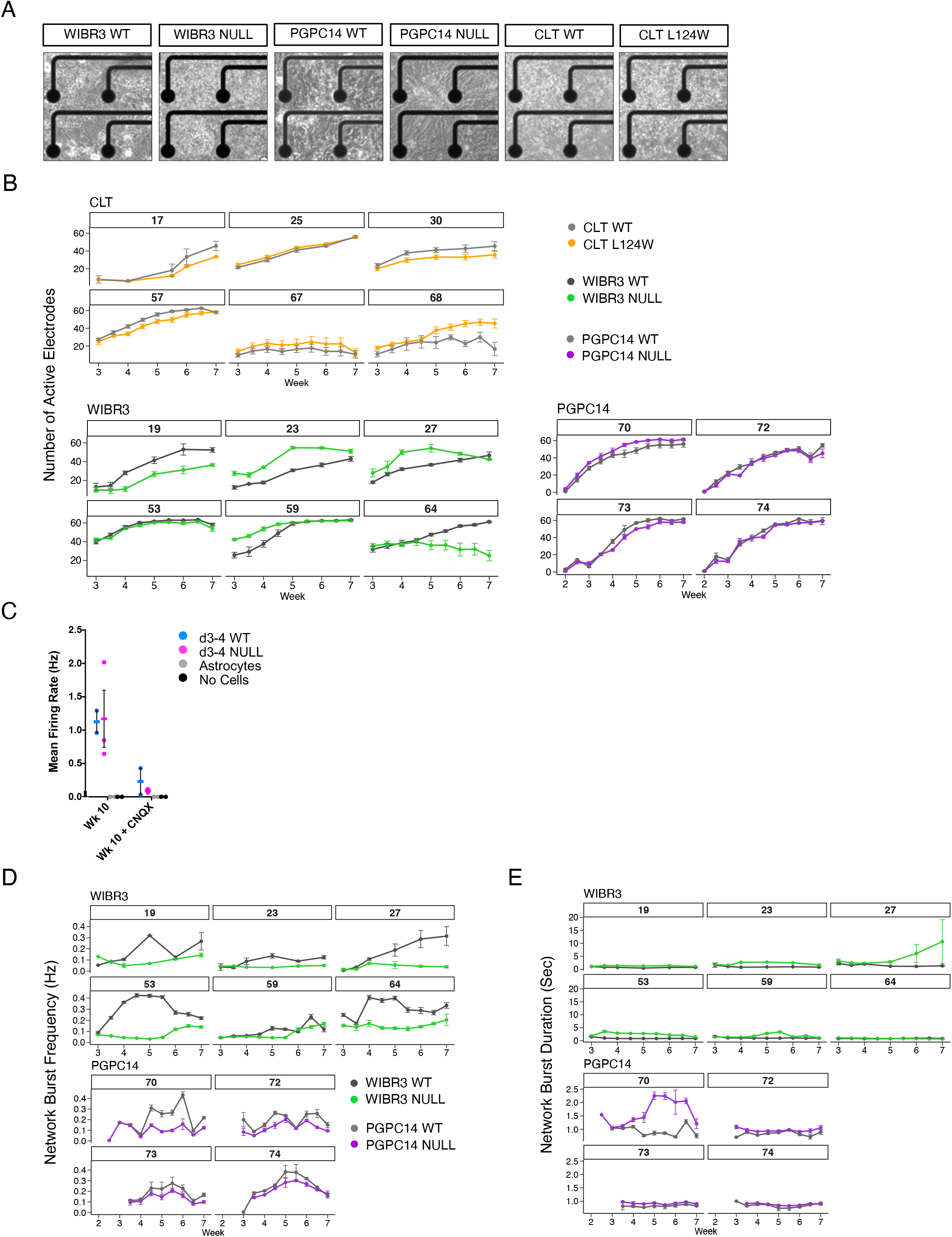
MEA properties in RTT Ngn2 neurons. **A.** Representative images of 5-7 week old cultures on MEA plates. **B.** Number of active electrodes for isogenic WT and RTT Ngn2 neurons over 3 to 7 weeks of individual replicate plates (number above indicate Plate ID). 6 replicate plates of CLT, 6 replicate plates of WIBR3 and 4 replicate plates of PGPC14 with 3-6 wells per genotype per plate. Mean +/- SEM shown. **C.** Mean firing rate of 10 week old WT and *MECP2* null *Ngn2* neurons treated with CNQX. **D-E.** Network burst frequency and duration for isogenic WT and null *Ngn2* neurons over 3 to 7 weeks of individual replicate plates (number above indicates Plate ID). 6 replicate plates of WIBR3 and 4 replicate plates of PGPC14. Mean with SEM.

**Figure S5.**
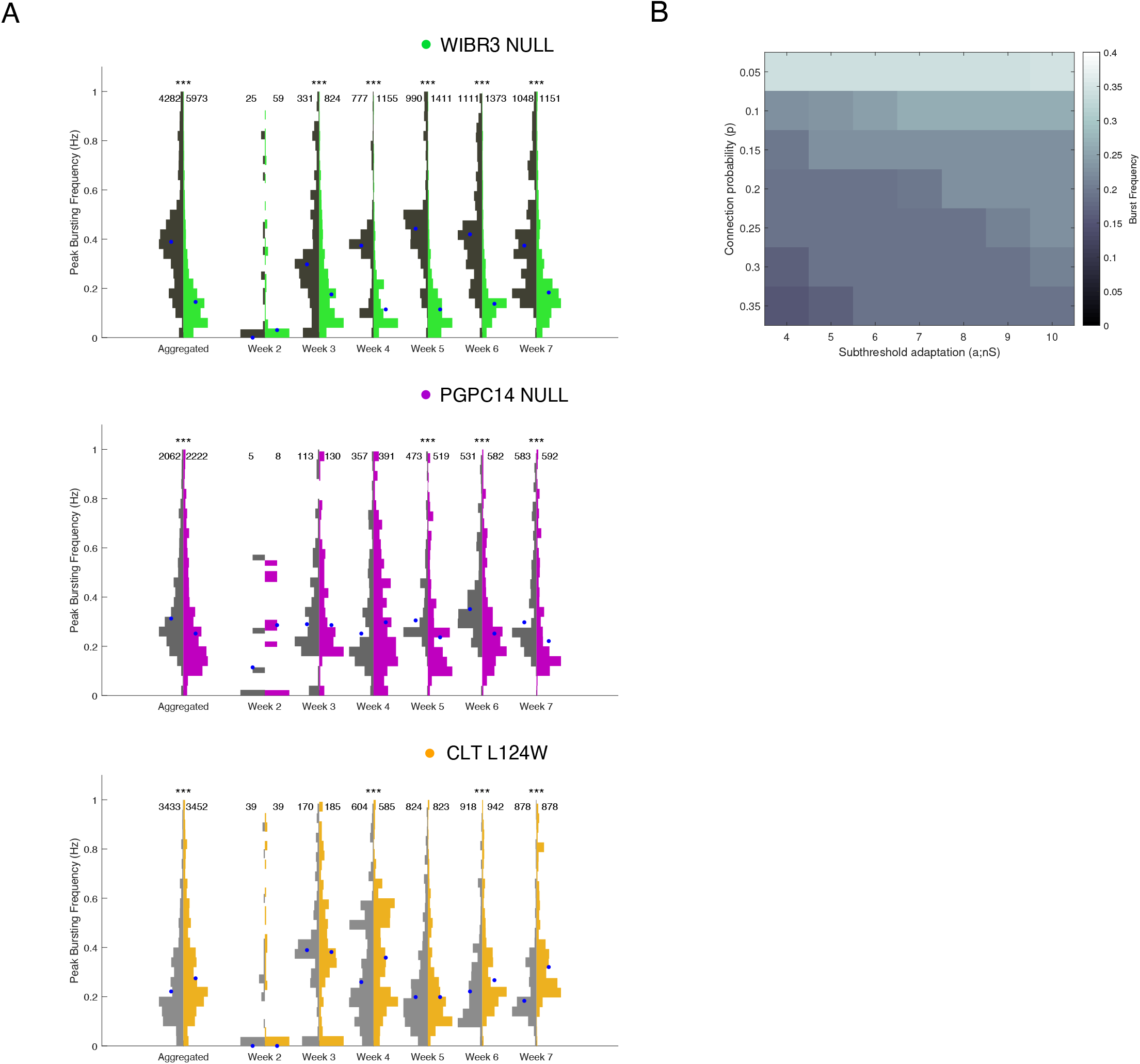
Network analysis in RTT Ngn2 neurons. **A.** Distribution of bursting frequencies across 6 WIBR3 plates, 4 PGPC14 plates, and 5 L124W CLT plates. Medians in each case are plotted as blue dots. Because only electrodes with detected bursts are plotted, the number of samples increases from week 2 to 7 (noted above each histogram). **B.** Control simulation to understand the effect of connectivity (vertical axis) and subthreshold adaptation (horizontal axis). Each point in the heatmap represents the measured burst frequency (colorbar) at one combination of connection probability and subthreshold adaptation. Statistical significance was evaluated by Mann Whitney test in Figure A. *p < 0.05, **p < 0.01, ***p < 0.001.

## Notes

### Summary of Updates

The revised manuscript includes a revised title, additional authors, updated abstract, updated and new text, updated figures and supplementary figures, and a new Figure 5 on simulation models.

